# Allostery is a widespread cause of loss-of-function variant pathogenicity

**DOI:** 10.1101/2025.06.20.660737

**Authors:** Xiaotian Liao, Ben Lehner

## Abstract

Allosteric communication between non-contacting sites in proteins plays a fundamental role in biological regulation and drug action. While allosteric gain-of-function variants are known drivers of oncogene activation, the broader importance of allostery in genetic disease and protein evolution is less clear. Here, we integrate large-scale experimental measurements and neural network models to provide evidence that allostery is a widespread cause of loss-of-function variant pathogenicity in human genetic diseases. In addition, our analyses reveal a conserved distance-dependent decay of allosteric mutational effects outside of protein active sites. As an important mechanism of pathogenicity, allostery needs to be better mapped, understood, and predicted across the human proteome.

## Introduction

Allostery, the regulation of a protein’s activity through changes at sites distant from an active site, underpins protein regulation^1–3^. Allostery allows proteins to function as molecular switches and microprocessors, with their enzymatic activities and molecular interactions quantitatively controlled by small molecule binding, post-translational modifications, and macromolecular interactions^4^. Allostery is central to the control of metabolic pathways, signal transduction, growth control and neural information processing in both normal physiology and disease^1,3,5–9^. Moreover, many effective therapeutics bind allosteric sites and have allosteric mechanisms-of-action, including drugs targeting ion channels, receptors, kinases, and transcription factors^2,10–14^.

The importance of allostery for human genetic disease is, however, less well established^6,15,16^. Allosteric gain-of-function (GOF) variants are a recognized class of activating mutations in human oncogenes^2,17^. For example, the G12C and other driver mutations in KRAS allosterically increase binding to effector proteins such as the RAF kinase^18,19^, and the V617F mutation within the pseudokinase domain of the JAK2 tyrosine kinase disrupts its autoinhibitory interaction with the kinase domain, leading to constitutive activation through trans-phosphorylation^20,21^. Similarly, allosteric mutations in the epidermal growth factor receptor (EGFR), including L858R and T790M, increase kinase activity and confer resistance to targeted therapies in non-small cell lung cancer^22,23^. An increasing number of licensed cancer therapeutics have allosteric mechanisms of action, including the first effective KRAS inhibitors^6,15,24^.

Most pathogenic variants causing human genetic diseases are, however, loss-of-function (LOF) mutations^25,26^. LOF variants include premature stop codons (PTCs) and splice-site variants, but also a large number of protein missense variants^27,28^. Missense variants that change the amino acid (AA) sequence of a protein can have diverse molecular effects. These include destabilization of a protein’s fold, with recent studies suggesting that 39-60% of clinically pathogenic missense variants reduce protein stability^29–33^. However, variants can also impair molecular activities beyond stability, such as disrupting molecular interactions or enzymatic activities^32,34–42^. We term these variants ‘functional mutations’ to distinguish them from variants affecting protein stability.

A small number of LOF variants are proposed to have allosteric mechanisms-of-action^6,25,43,44^. Examples include the G551D variant in the cystic fibrosis transmembrane conductance regulator (CFTR)^45–47^, as well as distal variants such as K246E and L311V in adenylosuccinate lyase (ADSL), which reduce enzymatic activity and cause ADSL deficiency—a metabolic disorder linked to autism and epilepsy^48^. Nevertheless, the broader contribution of allosteric loss-of-function variants to human genetic disease remains largely uninvestigated, a shortcoming that we seek to address in the present study.

Recent comprehensive mutagenesis studies have generated the first systematic experimental maps of allosteric communication throughout proteins^40–42,49,50^. These near complete atlases for a handful of proteins have revealed that many mutations—even those distant from active sites and binding sites—can exert measurable allosteric effects on protein function. For example, 114 amino acid substitutions in the oncoprotein KRAS allosterically inhibit binding to the oncoprotein effector RAF1^42^. Even in small protein domains such as SH3 domains, many variants have small but measurable allosteric effects^40^ that, in combination, can strongly impair binding to interaction partners^51^. These first allosteric maps suggest that allostery is widespread and highlight the need to reconsider the importance of allostery for variant pathogenicity and molecular evolution.

Here, we evaluate the contribution of allostery to loss-of-function human variant pathogenicity. Across extensive experimental datasets and state-of-the-art computational models, our analyses indicate that loss-of-function pathogenic variants frequently have allosteric mechanisms-of-action, disrupting protein function without destabilizing the fold or directly disrupting catalytic residues and binding sites. Across multiple analyses and structurally diverse proteins, these loss-of-function allosteric effects are enriched close to protein active sites and decay exponentially through protein structures. Our results indicate that allostery is a widespread and underappreciated mechanism of variant pathogenicity that needs to be better understood and predicted across the human proteome.

## Results

### Pathogenic variants often act through mechanisms beyond stability

From a structural perspective, missense mutations that impair protein function can act through three primary mechanisms: (1) globally destabilizing the protein fold, (2) directly disrupting the active or binding site, or (3) indirectly perturbing function through distal or allosteric effects. To estimate the proportion of mutations that act via destabilization, we first applied computational methods to predict the impact of each variant on protein stability. To assess the contribution of functionally disruptive mutations—those whose effects go beyond folding—we applied state-of-the-art deep learning approaches. Specifically, we used ThermoMPNN, an inverse folding neural network^52^ fine-tuned on large-scale experimental data^32^, to predict changes in folding free energy (ΔΔGf) for all possible missense mutations across 19,804 human proteins. This analysis encompassed 193,469,108 variants, including 113,994 annotated in the ClinVar database^53,54^ **(Fig. 1A, Supplemental Table 1)**.

**Fig. 1:**
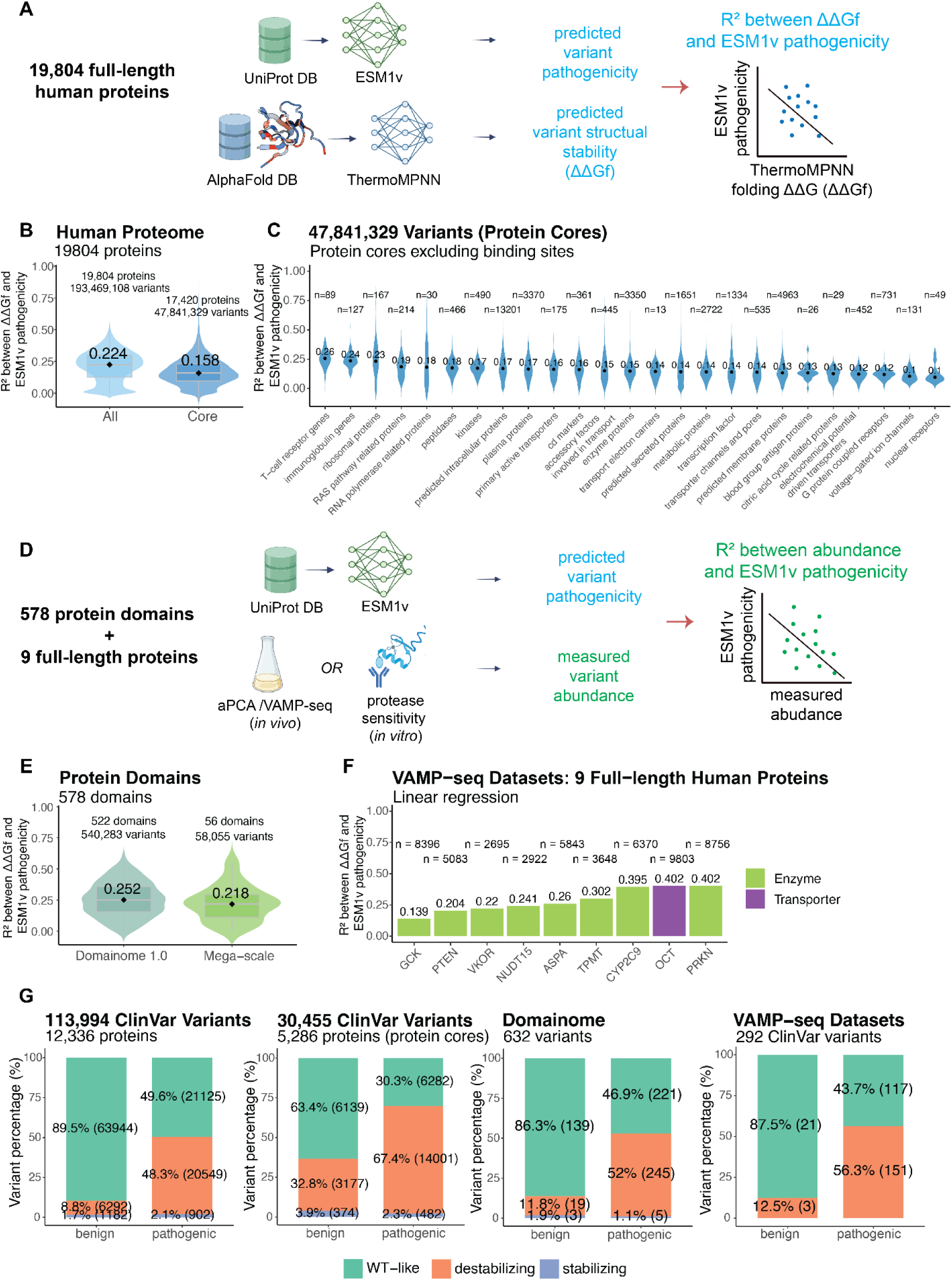
Pathogenic variants often act through mechanisms beyond stability. **A**. Overview of datasets used for proteome-wide computational analysis. Predicted pathogenicity scores were generated using ESM1v for all missense variants in 19,804 human proteins. Predicted folding stability changes (ΔΔGf) were computed using ThermoMPNN (kcal/mol). **B**. Distributions of R^2^ values from linear regression between predicted ΔΔGf and ESM1v scores for all missense variants across the human proteome (left), and for protein core variants (right), defined by excluding disordered, surface-exposed and predicted binding sites. Median R^2^ values are indicated. **C**. Distribution of R^2^ values, stratified by protein type (core variants only). Median R^2^ values are labeled. Protein types were defined based on the “Function and compartment based” classification provided by the Human Protein Atlas^83–85^. **D**. Overview of datasets used for large-scale experimental measurements. Protein abundance changes were quantified using multiple methods: *in vivo* DHFR complementation assay (Domainome 1.0^117^), *in vitro* cDNA display (Mega-scale^32^), and VAMP-seq^62^ (variant abundance by massively parallel sequencing) for full-length proteins. **E**. R^2^ distributions for abundance changes were regressed against ESM1v scores for the Domainome 1.0 and Mega-scale datasets. Median values are labeled. **F**. R^2^ values between ESM1v pathogenicity scores and variant-induced abundance changes for nine full-length proteins profiled using VAMP-seq. Protein types are indicated by bar color. The number of variants (n) and R^2^ values are shown. **G**. Classification of ClinVar missense variants into three categories: WT-like (green), destabilizing (orange), and stabilizing (blue), based on their impact on protein abundance. Percentages and total counts are shown for all ClinVar variants included in the proteome analysis (left), protein core variants (middle-left), variants from Domainome 1.0 (middle-right), and VAMP-seq (right), stratified by benign and pathogenic clinical labels.

To quantify how much of variant pathogenicity is attributable to changes in stability, we compared ThermoMPNN-predicted ΔΔGf values to ESM1v scores—a model that estimates the functional impact of mutations from evolutionary sequence conservation^55^. Across the entire proteome, predicted ΔΔGf explained only a modest proportion of the variance in ESM1v pathogenicity scores **(Fig. 1B, median R^2^ = 0.224, interquartile range (IQR) 0.125-0.298)**, suggesting that folding stability is only one of several mechanisms underlying missense variant pathogenicity.

Since most active and binding residues are solvent-exposed, we reasoned that analyses limited to buried residues would isolate stability-driven effects. However, even when excluding solvent exposed residues (relative solvent accessible surface area, rSASA > 0.2), predicted changes in stability still only partially explained predicted pathogenicity (Fig**. 1B****, median R^2^ = 0.158, IQR 0.096 - 0.232)**. This lower R^2^ in buried regions may in part reflect a more restricted range of ESM1v scores in the protein core, limiting the total explainable variance. The pattern held across functional classes: folding stability accounted for just ∼10% of pathogenicity in voltage-gated ion channels and nuclear receptors, while explaining ∼23-26% in ribosomal proteins, immunoglobulins, and T-cell receptors **(Fig. 1C)**.

Thus, state-of-the-art models suggest that stability is an important component of variant pathogenicity, but they predict that many variants are detrimental because of functional effects beyond stability changes.

#### Analysis of large-scale experimental abundance datasets

To go beyond computational predictions, we next turned to direct experimental measurements of changes in stability. We analysed data from two recent large-scale studies that together experimentally quantified changes in abundance for 598,338 variants in 578 different protein domains^32,33^. The first ‘MegaScale’ dataset used protease-sensitivity to quantify changes in folding free energy for 272,712 missense variants in 331 protein domains *in vitro*. The second ‘Domainome’ dataset used a protein-fragment complementation assay to quantify changes in cellular abundance for 607,081 missense variants in 522 protein domains **(Fig 1D)**. For variants with matching ESM1v scores, the experimentally-quantified abundance changes still only partly account for ESM1v-predicted pathogenicity **(Fig. 1E, median R^2^ = 0.252 and 0.218, respectively)**. These results match the computational predictions: many variants appear pathogenic despite maintaining near-normal stability or expression.

To assess whether this trend holds in full-length, physiologically relevant contexts, we analysed experimental abundance measurements for nine complete human proteins (Variant Abundance by Massively Parallel sequencing - ‘VAMP-seq’ dataset), spanning various functions and sizes **(Supplemental Table 2)**. Across these full-length proteins, abundance changes explained a median of only 26% of ESM1v variance **(Fig. 1F, R^2^ range: 13.9% – 40.2%)**, closely matching the domain-level findings.

#### Evaluation of ClinVar variants

Although protein language models provide a useful proxy for pathogenicity, we sought to validate our findings using clinically annotated variants. To this end, we turned to ClinVar, a curated database that classifies variants as pathogenic, benign, or of uncertain significance^53,54^. Comparing 42,576 pathogenic variants against 71,418 benign variants, we found that pathogenic variants are far more likely to be predicted as destabilizing **(Fig. 1G, 48.3% vs 8.8% odds ratio (OR) = 9.653, p < 2.2e-16, Fisher’s exact test (FET))**. Predicted stability changes are thus a very useful, albeit imperfect, predictor of pathogenicity (Receiver Operating Characteristic – Area Under the Curve (ROC-AUC) = 0.697, Precision–Recall Area Under the Curve (PR-AUC) = 0.618, n = 42,576 pathogenic, n = 71,418 benign variants).

As expected^56^, mutations in the buried protein cores were more likely to be destabilizing (OR = 8.072, p < 2.2e-16, FET) and also more likely to be pathogenic (OR = 4.243, p < 2.2e-16, FET). In total, 67.4% of pathogenic variants in the core were predicted to be destabilizing compared to 32.8% of benign core variants **(Fig. 1G)**. Variations were observed across protein functional classes, with the proportion of stability-affecting (destabilizing or stabilizing) pathogenic variants ranging from 38-43% in secreted proteins, voltage-gated ion channels, and transcription factors to 60% in metabolic proteins **(Fig. S2A)**.

Experimentally-measured stability changes presented a very similar picture: 52% of clinically pathogenic variants in protein domains and 56.3% of pathogenic variants in full-length proteins are measured to disrupt stability **(Fig. 1G)**. Thus, while experimentally-measured stability changes are predictive of pathogenicity—ROC-AUC = 0.70 and PR-AUC = 0.87 for domains (n = 471 pathogenic, 161 benign); ROC-AUC = 0.72 and PR-AUC = 0.96 for full-length proteins (n = 268 pathogenic, 24 benign)—additional pathogenic mechanisms must be widespread.

Together, these data demonstrate that while loss of stability is a common mechanism of pathogenicity, a substantial fraction of pathogenic variants must act through other mechanisms—such as disrupting active sites, perturbing binding interfaces, or exerting long-range allosteric effects.

#### Functional effects decay exponentially with distance

Allosteric propagation—the transmission of energetic changes from one site in a protein to another—provides a mechanistic basis for how mutations distant from active sites can still impact function^57,58^. As a baseline, in the absence of allosteric effects, one would expect no relationship between functional effect size and distance from active site. Mutations would primarily act locally, and effects would cluster strictly at active sites. However, in the presence of allostery, mutational effects are expected to propagate from active sites through the structure, and we would expect functional effects to decay with 3D distance—strongest near the active site and weaker further away. This distance-dependent allosteric decay is visible in the small number of experimental allosteric maps generated to date^40–42,49,50^.

To test whether variant effects beyond stability changes include long-range allosteric mechanisms, we first analysed missense mutations across six structurally and functionally diverse full-length human allosteric proteins **(Fig. 2A–B, Supplemental Table 3)**. For each allosteric protein in our test set, we compared predicted pathogenicity (ESM1v) to predicted folding stability changes (ΔΔGf) from ThermoMPNN. We then fitted a Locally Estimated Scatterplot Smoothing (LOESS) curve and computed residuals to quantify variant effects not attributable to stability changes **(Fig. 2C, Fig. S3B)**.

**Fig. 2:**
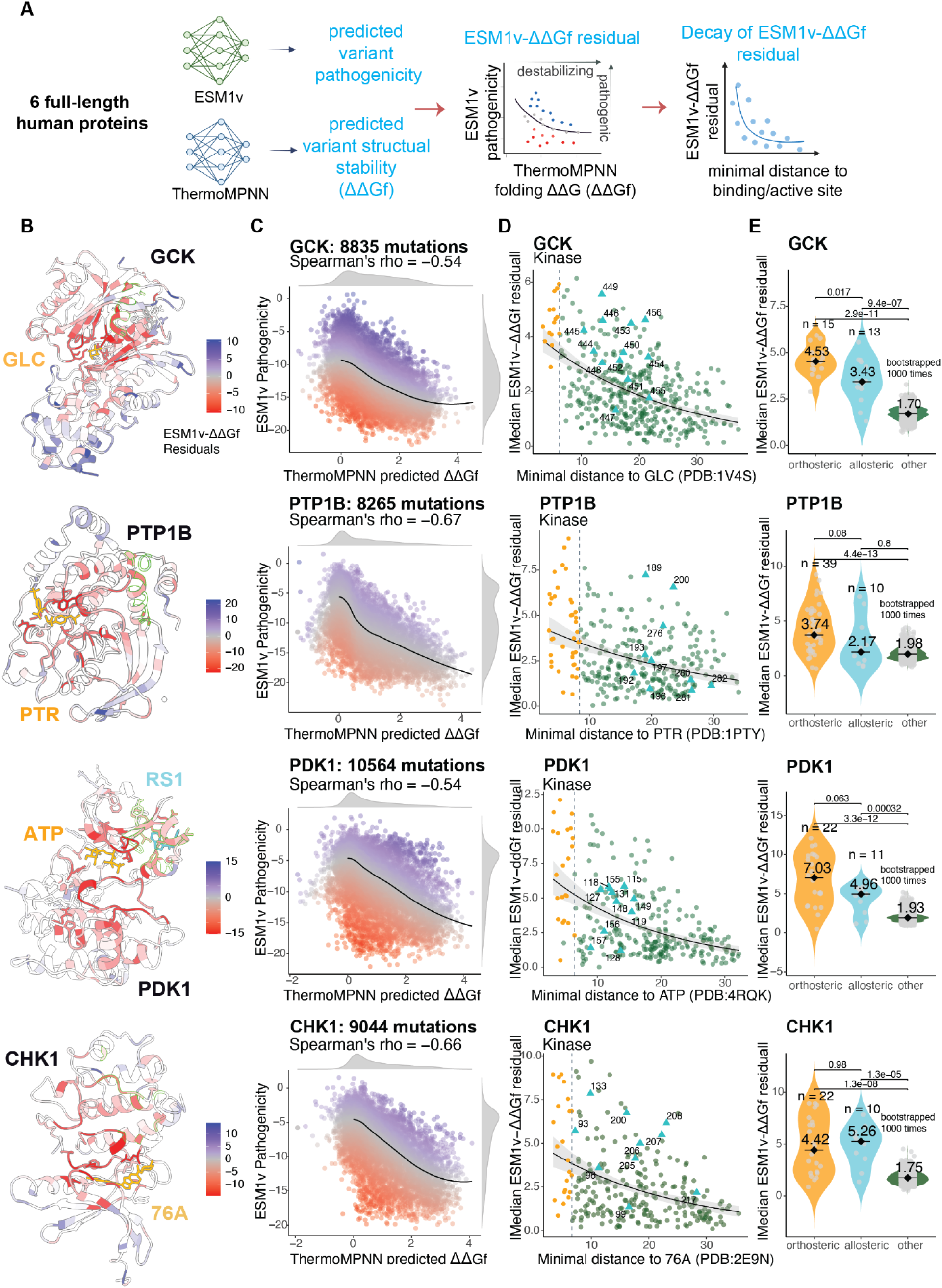
Functional effects of mutations decay exponentially with distance. **A**. Overview of the analytical framework. Predicted variant pathogenicity scores from ESM1v and folding stability predictions from ThermoMPNN (ΔΔGf) were used to compute residuals (ESM1v–ΔΔGf), representing functional effects not explained by structural destabilization. These residuals were then analyzed for spatial decay relative to distance from the substrates. **B**. Crystal structures of GCK (PDB: 1V4S^69^), PTP1B (PDB: 1PTY^118^), PDK1 (PDB: 4RQK^59^), and CHK1 (PDB: 2E9N^119^), colored by the median residuals between ESM1v and ThermoMPNN (TMPNN) predicted folding free energy changes (ΔΔGf) values. Bound substrates — glucose (GLC), O-phosphotyrosine (PTR), adenosine triphosphate (ATP), and CHK1 inhibitor (76A) — are shown in orange. Allosteric inhibitor RS1 is labeled in blue. Annotated allosteric positions are highlighted in neon green. **C**. Scatter plots of ThermoMPNN-predicted ΔΔGf versus ESM1v scores for missense mutations in each protein. A LOESS curve (black line) models the trend, with residuals quantifying deviation between folding impact and predicted pathogenicity. Stable mutations cluster around ΔΔGf ≈ 0. **D**. Per-residue relationship between the absolute median ESM1v–ΔΔGf residual and the minimal Cα distance (Å) to the substrates for each structure. Residuals are calculated using only mutations in non-orthosteric sites with negative residuals. An exponential decay model (black curve) was fitted to the data, with 95% confidence intervals shown as grey shading. The grey dashed line indicates the maximum of the minimal distances from any substrate-contacting (orthosteric) residue. Sites are colored by functional class: orthosteric sites in orange circles, non-orthosteric sites in dark green circles, and known allosteric sites (i.e., sites in physical contact with known allosteric regulators) in cyan triangles. **E**. Distributions of per-site absolute median residuals between ESM1v and ΔΔGf for orthosteric, allosteric, and other sites. Each point represents the absolute median ESM1v-ΔΔGf residual for a site. For orthosteric and allosteric sites, residuals are calculated directly per annotated site. For the ‘other’ group, distributions are derived from 1,000 bootstrap replicates, each sampling the same number of non-orthosteric and non-allosteric sites as in the orthosteric group. Violin plots show the distribution shape; black bars and diamonds indicate medians. Sample sizes and median values are annotated. Statistical significance between groups was assessed using two-sided Wilcoxon rank-sum tests.

Across all six proteins, many variants exhibited large residuals, indicating functional effects not accounted for by changes in fold stability. ESM1v and ΔΔGf were moderately correlated (Spearman’s ρ = –0.25 to –0.67), with enrichment of large residuals at annotated orthosteric sites and distal regulatory positions where known allosteric regulators bind **(Fig. 2B, D, Fig. S3A, C)**.

We next asked whether these stability-independent functional effects follow a spatial pattern consistent with allosteric propagation^40–42,49,57,58^. For each residue, we computed its minimal distance to the ligand atoms and plotted the absolute median residual against this distance. In all proteins, the absolute median residuals were strongest for residues close to the active site, with the predicted functional effects of mutations decaying approximately exponentially with 3D distance from the ligand **(Fig. 2D, Fig. S3B)**. This trend was statistically significant: in a linear model regressing the log-transformed absolute residuals on distance, increasing distance was significantly associated with weaker functional effects **(Supplemental Table 3)**. Across proteins, the half-distances for decay (d_1/2_) are comparable to the distance-dependent decay of allosteric mutational effects observed in experimental allosteric maps **(GCK: 15.7 Å, PTP1B: 19.7Å, PDK1: 12.3Å, CHK1: 15.9 Å; Supplemental Table 3)**^40–42,49,50^.

To determine whether the absolute median residuals are increased at known allosteric sites, we compared their magnitudes at orthosteric sites, distal sites where known allosteric regulators bind, and randomly sampled control positions matched for protein abundance. In four out of the six proteins (all except PTP1B and CASP1), absolute median residuals were significantly higher at both orthosteric and known allosteric sites than at control positions **(Fig. 2E, Fig. S3D)**. In PTP1B and CASP1, allosteric sites also showed larger absolute residuals on average **(Fig. S3E)**, although the difference was not statistically significant. It is important to note that this definition of allosteric sites is likely conservative, as it is based only on the binding sites of known allosteric regulators. Most allosteric proteins will harbour additional allosteric sites that remain untested.

Notably, the distance-dependent decay is not uniform. Some positions exhibit residuals much larger than expected for their distance from the active site. We refer to these as allosteric hotspots—residues with more functional impacts than expected. These hotspots include binding sites of known allosteric regulators, such as inhibitors of protein tyrosine phosphatase 1B (PTP1B) and compound RS1, which targets pyruvate dehydrogenase kinase 1 (PDK1).^59,60^.

These results suggest that allosteric mechanisms potentially underlie a substantial and underappreciated component of pathogenicity in missense variants, and that the magnitude of these effects decays in a conserved, distance-dependent manner from functional sites. To benchmark the validity of our computational predictions, we compared ESM1v–ThermoMPNN ΔΔGf residuals to experimentally measured, stability-independent activity effects from published deep mutational scanning maps^38,40,61–63^. Across all proteins for which matched in silico and experimental data were available, we observed a consistent and significant correlation between computational residuals and experimentally defined allosteric effects **(Fig. S4)**, supporting the relevance of our approach.

### Experimental allosteric maps confirm distance-dependent decay

While computational predictions enable proteome-wide predictions of allosteric variant effects, they do not directly measure functional consequences of mutations. To address this limitation, we turned to experimental allosteric maps that quantify the impact of every missense mutation on both folding and binding—across a smaller but well-characterized set of proteins.

Recent advances have enabled the generation of comprehensive experimental allosteric maps for individual proteins and protein domains^40–42,49,50^. For protein–protein interaction domains, these maps report the effect of every possible missense mutation on folding free energy (ΔΔGf) and binding free energy (ΔΔGb), allowing measurement of both stability and functional consequences of mutations at each site^40,42^. In these systems, mutations that alter binding without directly contacting the ligand must, by definition, act through an allosteric mechanism. To evaluate this, we focused on two domains with complete mutational maps: the third PDZ domain of PSD-95 and the SH3 domain of GRB2 **(Fig. 3A)**.

**Fig. 3:**
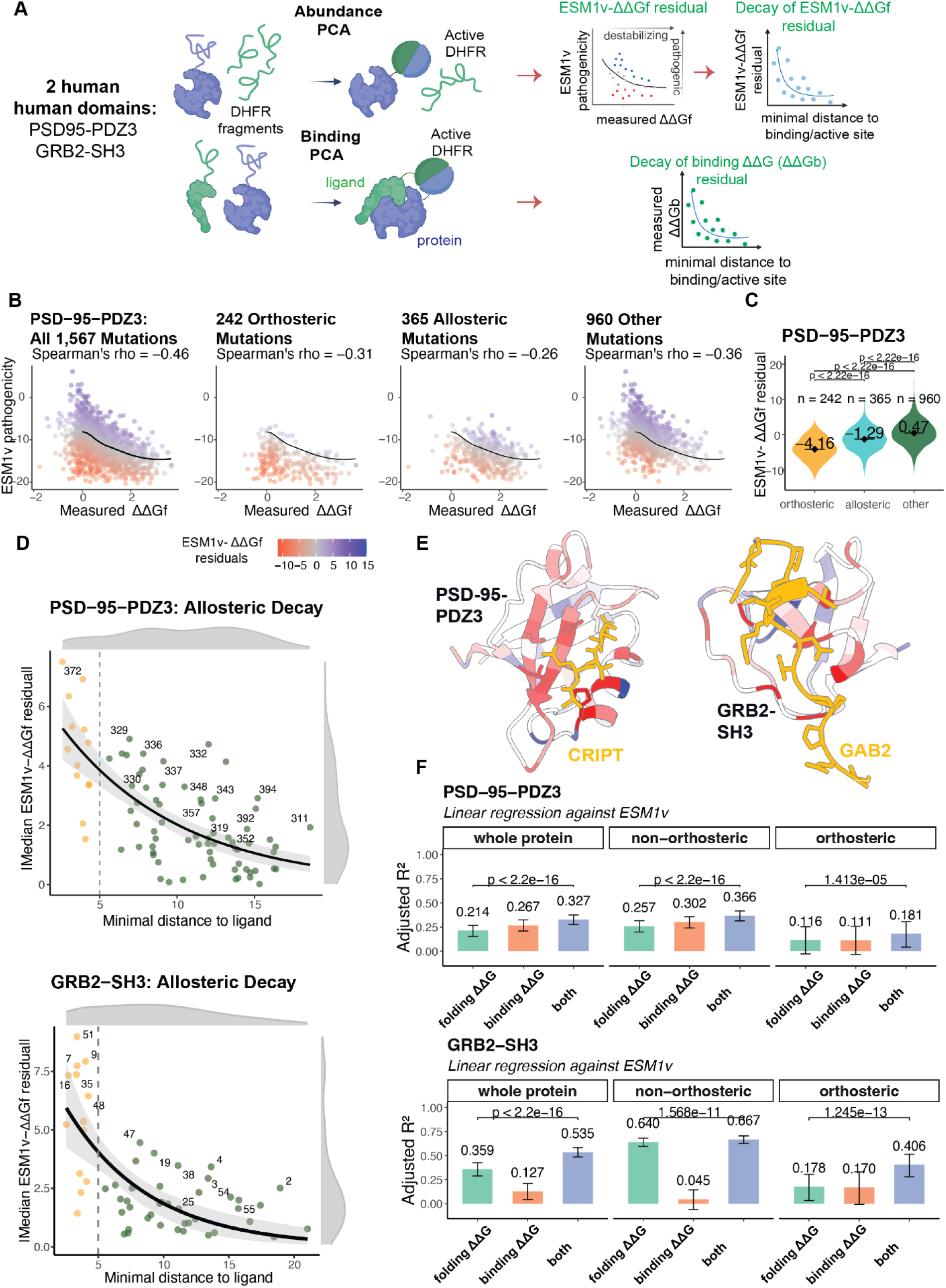
Experimental allosteric maps confirm distance-dependent decay of functional mutational effects. **A**. Overview of the functional assays used to assess the effects of missense mutations on protein abundance and binding in two human domains: PSD-95-PDZ3 and GRB2-SH3. Protein abundance and binding were quantified using protein fragment complementation (PCA) assays (abundance PCA: aPCA, binding PCA: bPCA). The residual between ESM1v scores and experimentally measured folding free energy changes (ΔΔGf) captures functional effects not explained by stability and is used to assess distance-dependent allosteric decay. **B**. Residual plots showing the relationship between ESM1v scores and experimentally measured ΔΔGf values for all PSD-95 PDZ3 mutations (left), orthosteric mutations (middle left), allosteric mutations (middle right), and all other mutations (right). LOESS regression lines (black) show the trend across all non-orthosteric mutations. **C**. Distributions of the ESM1v–ΔΔGf residuals per variant group. Significance values were calculated using the two-sided Wilcoxon rank-sum test. **D**. Relationship between the absolute median ESM1v–ΔΔGf residual and the minimal side chain distance to the ligand in PSD-95 PDZ3 (top) and GRB2 SH3 (bottom). Residuals were calculated from mutations with negative values only. An exponential decay model (black curve) was fitted to the data, with 95% bootstrap confidence intervals shown as grey shading. The grey dashed line marks the maximal minimal distance of orthosteric residues (i.e. the outer bound of direct ligand contacts). Residues are color-coded by functional class: orthosteric (orange) and non-orthosteric (dark green). Residues with significantly greater residuals than expected under the decay model (FDR-adjusted *p* < 0.05, one-sided Wilcoxon signed-rank test) are labeled by position number. **E**. Crystal structures of PSD-95 PDZ3 bound to CRIPT (PDB: 1BE9^120^) and GRB2-SH3 bound to GAB2 (PDB: 2VWF^121^), colored by median ESM1v–ΔΔGf residuals for each site. **F**. Linear regression models quantifying the variance in ESM1v scores explained by measured ΔΔGf (folding free energy changes), ΔΔGb (binding free energy changes), or both, across orthosteric and non-orthosteric sites. Adjusted R^2^ values are shown with bootstrapped confidence intervals. Statistical significance was evaluated using ANOVA to compare each baseline model to the corresponding full model including both energy terms.

As observed in our proteome-wide analyses, folding stability changes alone explained only a modest fraction of predicted pathogenicity: R^2^ = 0.214 for PSD-95 and R^2^ = 0.359 for GRB2 **(Fig. 3F)**. To isolate the stability-independent component of pathogenicity, we fitted a LOESS curve between ΔΔGf and ESM1v predicted pathogenicity for each domain and computed residuals, quantifying the extent to which mutations disrupt function beyond what would be expected from their impact on folding stability alone **(Fig. 3B, Fig. S5A)**.

As expected, the largest absolute median residuals were observed at orthosteric binding residues and at experimentally annotated allosteric mutations **(Fig. 3C, Fig. S5B)**. These effects also followed a spatial pattern consistent with allosteric propagation: absolute residual magnitudes were highest near the binding interface and decayed with increasing 3D distance from the ligand **(Fig. 3D, PSD-95-PDZ3: slope = -0.14, *P* = 2.46e-07; GRB2-SH2: slope = -0.09, *P* = 2.06e-05)**. Visualizing the median ESM1v-ΔΔGf residual per site further illustrates this distance dependence of predicted functional effects away from the binding interface **(Fig. 3E)**.

We next evaluated experimentally measured ΔΔGb values to directly assess allosteric effects. For residues outside the binding interface, changes in ΔΔGb isolate the allosteric component of binding disruption. These values explained 30.2% of ESM1v variance in PSD-95 and 4.5% in GRB2 **(Fig. 3F)**. The smaller contribution of ΔΔGb to pathogenicity in GRB2-SH3 likely reflects the domain’s very small size and reduced allostery.

Importantly, binding free energy changes (ΔΔGb) also showed a distance-dependent decay from the ligand **(Fig. S5C)**. Across residues, ΔΔGb values correlated with the magnitude of ESM1v–ΔΔGf residuals **(Fig. S5D)**, with a stronger relationship in GRB2 (Spearman’s ρ = –0.70) than in PSD-95 (ρ = –0.31). Moreover, models incorporating both ΔΔGf and ΔΔGb explained more pathogenicity variance than models using stability alone—even after excluding mutations within the binding interface **(Supplemental Tables 4–7)**. This further demonstrates the allosteric contribution to pathogenicity is independent of changes in fold stability.

### Clinical pathogenic variants in PTEN and GCK show allosteric decay

Finally, to further evaluate the contribution of allostery to clinical pathogenicity, we analysed two human proteins with many known disease-causing variants and large-scale experimental data: the lipid phosphatase PTEN and the glucose-sensing kinase glucokinase (GCK) **(Fig. 4A, 5A)**. For both proteins, the effects of thousands of variants on abundance and enzymatic activity had been measured, enabling direct assessment of functional impact—including effects that are independent of abundance.

**Fig. 4:**
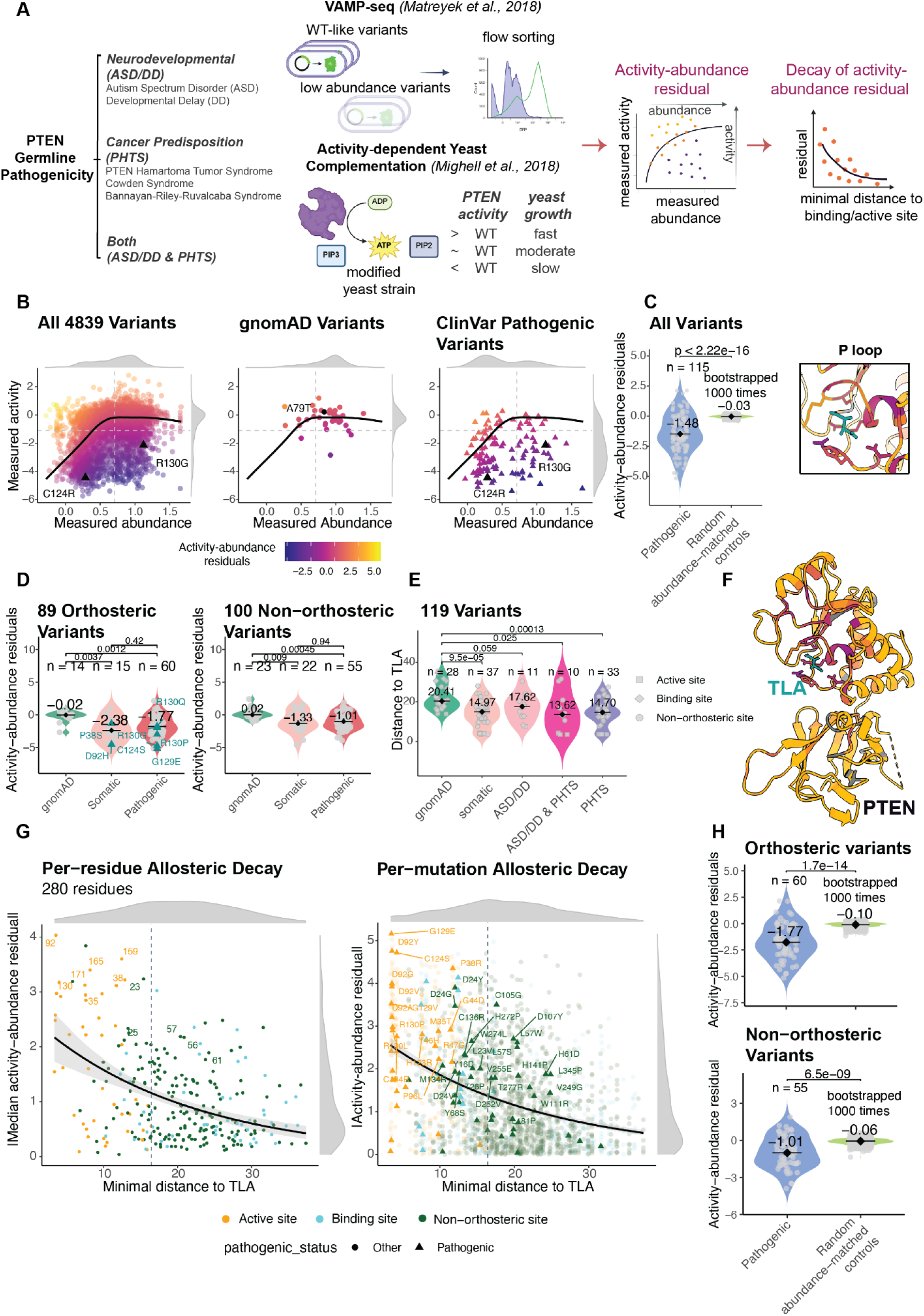
Clinically pathogenic variants in PTEN show allosteric decay. **A**. Overview of the functional assays used to measure the effects of PTEN missense mutations on protein abundance and activity. Protein abundance was assessed using VAMP-seq, and phosphatase activity using activity-dependent yeast complementation. The residual between measured activity and abundance captures non-stability-mediated functional changes, and is used to assess distance-dependent allosteric decay. **B**. Scatter plots of measured abundance versus activity fitness for all PTEN variants (left), gnomAD variants (middle), and ClinVar pathogenic variants (right). A LOESS regression curve (black line) models the relationship. Residuals from this fit quantify deviations from the abundance-activity trend. Pathogenic variants are marked with triangles, and a benign variant with a black circle. Dashed grey lines indicate wild-type abundance (0.71) and activity (-1.11) values, as determined in previous publications^61,62,72^. Two canonical pathogenic mutations at active sites, C124R and R130G, are highlighted with black triangles. **C**. Distributions of activity–abundance residuals for pathogenic and control variants. For each group, residuals reflect the deviation between observed activity and that predicted from abundance. Pathogenic variant residuals are compared to a bootstrapped null distribution generated from 1,000 abundance-matched samples of non-pathogenic variants (matching within 0.5-unit abundance bins). Violin plots show the distribution of median residuals, with medians indicated by black bars and diamonds. Sample sizes and median values are annotated. Statistical significance between groups was assessed using two-sided Wilcoxon rank-sum tests. **D**. Violin plots comparing activity-abundance residuals across variant sources (gnomAD, somatic, ClinVar pathogenic), separated into orthosteric (active/binding site) and non-orthosteric variants. Known dominant negative variants are labeled in cyan. **E**. Violin plots showing the minimal Cα distance of each variant to TLA across variant classes. **F**. Crystal structure of PTEN (PDB: 1D5R^102^) colored by the median activity-abundance residual at each position. L(+)-tartaric acid (TLA), located near the catalytic site, is shown in cyan. A close-up of the P loop highlights catalytic residues with large residuals. **G**. **Left:** per-residue relationship between absolute median activity-abundance residual and minimal distance to TLA for all 280 PTEN residues. An exponential decay model (black curve) was fit to the data, with 95% confidence intervals shown as grey shading. The vertical dashed line marks the maximum distance among annotated active site residues. Residues with significantly stronger effects than expected under the decay model (FDR-adjusted *p* < 0.05, one-sided Wilcoxon test) are labeled. **Right:** per-mutation relationship between absolute activity-abundance residual and distance for all variants with negative residuals. An exponential decay model (black curve) was fitted to the data, and 95% confidence intervals from bootstrapping are shown as grey shading. The dashed grey line indicates the maximal distance from any active-site residue to the ligand. Each point represents a single variant, colored by site type—active site (orange), membrane or calcium-binding site (cyan), or non-orthosteric site (dark green)—and triangle shaped/alpha-scaled by ClinVar pathogenicity classification. Labels mark pathogenic mutations at active or non-orthosteric sites with residuals that lie above the 95% confidence interval of the decay model, indicating stronger-than-expected divergence from abundance-predicted activity. **H**. Distributions of activity–abundance residuals for pathogenic and control variants, stratified by whether variants are orthosteric or non-orthosteric. Statistical significance between groups was assessed using two-sided Wilcoxon rank-sum tests.

PTEN germline variants are associated with a spectrum of disorders, including macrocephaly, autism spectrum disorder (ASD), developmental delay, and PTEN hamartoma tumor syndrome (PHTS), which encompasses Cowden syndrome and other cancer predisposition syndromes^64^.

Based on established criteria^65^, we grouped PTEN variants into three phenotypic categories: (1) neurodevelopmental, (2) cancer-predisposition, and (3) combined. In addition to these inherited phenotypes, PTEN is a well-established tumour suppressor, with somatic missense mutations frequently acting as drivers in glioblastoma, endometrial, prostate, and breast cancers^66^.

GCK germline mutations cause two opposing metabolic disorders depending on how they affect enzymatic function. Loss-of-function variants impair glucose sensing and lead to GCK-MODY (Maturity-Onset Diabetes of the Young, type 2), a mild familial form of hyperglycemia. In contrast, gain-of-function mutations cause congenital hyperinsulinemic hypoglycemia (HH), a neonatal disorder of excessive insulin secretion and persistent hypoglycemia^63,67,68^.

In PTEN, intracellular abundance was measured for 5,083 missense variants using VAMP-seq, and phosphatase activity was measured for 7,260 variants via a yeast-based complementation assay^61,62^ **(Fig. 4A)**. In GCK, abundance (8,396 variants) and catalytic activity (8,570 variants) were similarly quantified using yeast-based systems^38,63^ **(Fig. 5A)**.

**Fig. 5:**
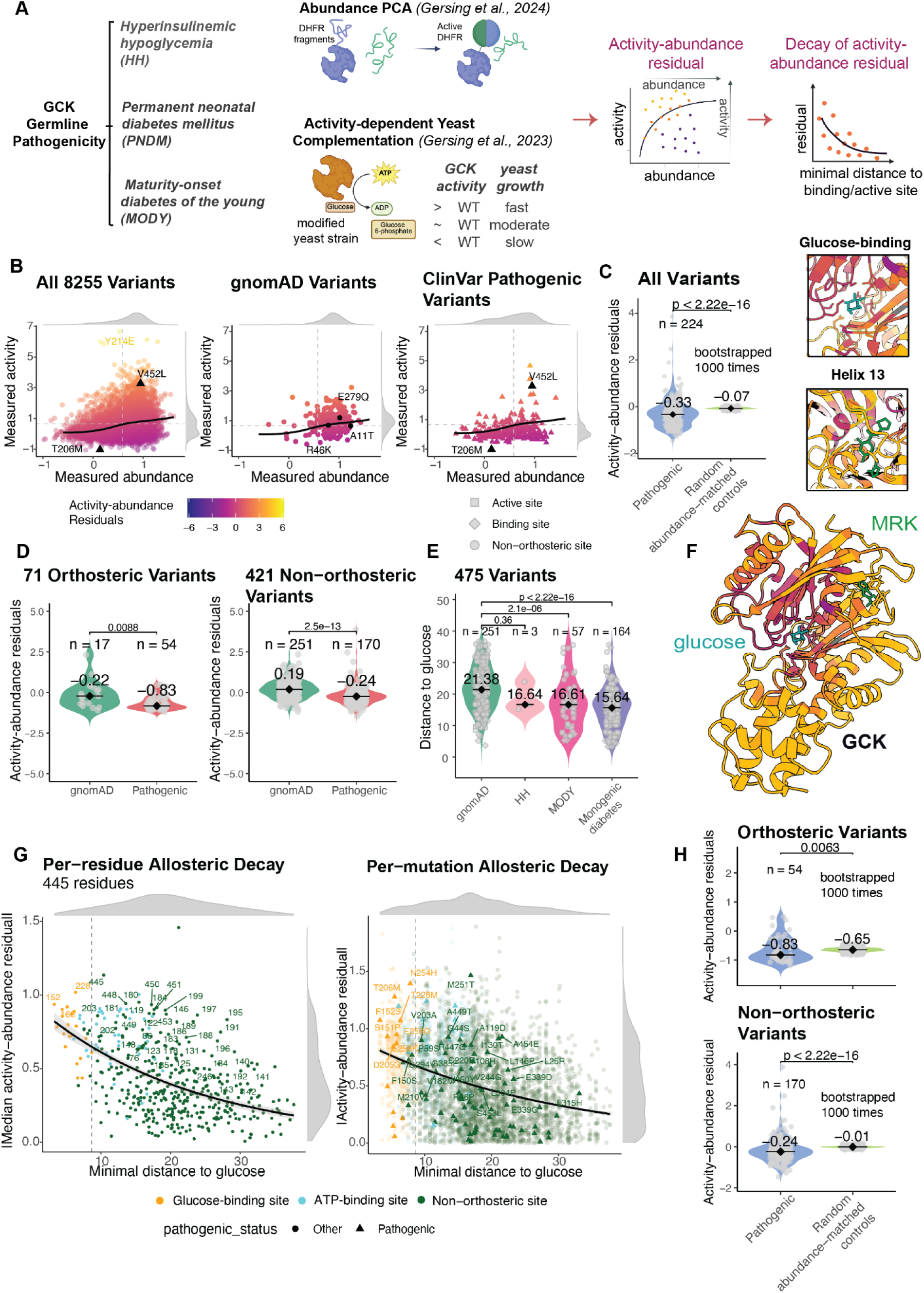
Clinically pathogenic variants in GCK show allosteric decay. **A**. Overview of the functional assays used to assess the effects of GCK missense mutations on protein abundance and activity. Protein abundance was measured using abundance PCA, and glucokinase activity was measured via a yeast complementation system. The residual between activity and abundance represents non-stability-mediated functional disruption and is used to assess distance-dependent allosteric effects. **B**. Scatter plots of measured abundance versus activity fitness scores for all GCK variants (left), gnomAD variants (middle), and ClinVar pathogenic variants (right). A LOESS regression curve (black line) models the overall trend. Residuals from this fit capture deviations between abundance and activity. ClinVar pathogenic variants are shown as triangles, and benign variants as circles. Grey dashed lines indicate wild-type abundance (0.58) and activity (0.66) values, as determined in previous publications^38,63^. Canonical gain-of-function pathogenic variant (V452L) and loss-of-function pathogenic variant (T206M) are noted in black triangles. **C**. Distributions of activity–abundance residuals for pathogenic and control variants. For each group, residuals reflect the deviation between observed activity and that predicted from abundance. Pathogenic variant residuals are compared to a bootstrapped null distribution generated from 1,000 abundance-matched samples of non-pathogenic variants (matching within 0.5-unit abundance bins). Violin plots show the distribution of median residuals, with medians indicated by black bars and diamonds. Sample sizes and median values are annotated. Statistical significance between groups was assessed using two-sided Wilcoxon rank-sum tests. **D**. Violin plots comparing activity-abundance residuals across gnomAD and ClinVar pathogenic variants, separated by whether the variant lies in orthosteric (ATP/glucose-binding) or non-orthosteric sites. **E**. Violin plots comparing the minimal Cα distance to glucose across gnomAD and different ClinVar pathogenic variant classes (HH, MODY, monogenic diabetes). **F**. Structure of GCK (PDB: 1V4S) colored by median activity-abundance residual per residue. Glucose is labeled in cyan. Key functional regions, including glucose-binding sites and helix 13 (residues 444–456), are highlighted in close-up views. **G**. **Left:** per-residue analysis showing the relationship between absolute median activity-abundance residual and minimal distance to glucose across 445 GCK residues. An exponential decay model (black curve) was fit to the data, with 95% confidence intervals shown as grey shading. The vertical dashed line marks the maximum distance among annotated active site residues. Residues with significantly stronger effects than expected under the decay model (FDR-adjusted *p* < 0.05, one-sided Wilcoxon test) are labeled. **Right:** per-mutation analysis showing absolute residuals versus distance for mutations with negative residuals. An exponential decay model (black curve) was fitted to the data, and 95% confidence intervals from bootstrapping are shown as grey shading. The dashed grey line indicates the maximal distance from any glucose-binding site residue to the ligand. Each point represents a single variant, colored by site type—glucose-binding site (orange), ATP-binding site (cyan), or non-orthosteric site (dark green)—and triangle shaped/alpha-scaled by ClinVar pathogenicity classification. Labels mark pathogenic mutations at glucose-binding or non-orthosteric sites with residuals that lie above the 95% confidence interval of the decay model, indicating stronger-than-expected divergence from abundance-predicted activity. **H**. Distributions of activity–abundance residuals for pathogenic and control variants, stratified by whether variants are orthosteric or non-orthosteric. Statistical significance between groups was assessed using two-sided Wilcoxon rank-sum tests.

To isolate functional effects beyond abundance changes, we fitted LOESS curves between abundance and activity using only non-orthosteric variants to define a baseline relationship. Residuals from this fit capture changes in activity not explained by changes in protein abundance **(Fig. 4B, 5B)**.

As expected, many pathogenic ClinVar variants reduced protein abundance. However, many others reduced enzymatic activity more than expected given their abundance levels **(Fig. 4C, 5C)**. Mapping these residuals onto the protein structures revealed their enrichment in both active sites and distal allosteric regions. In PTEN, high-residual variants localized to the catalytic core and surface-exposed residues, including the WPD, P, and TI loops that form the active site pocket **(Fig. 4F)**. In GCK, pathogenic variants clustered not only in the glucose-binding pocket but also in helix 13, a known allosteric regulator of activity^69,70^ **(Fig.5F)**.

In contrast, common variants from gnomAD and clinically benign variants (e.g., PTEN A79T; GCK A11T, R46T, E279Q) generally maintained wild-type-like abundance and activity, and showed minimal residuals^71^ **(Fig. 4B,D**; **Fig. 5B,D)**. Among pathogenic variants, the activity residuals were often especially large for variants that did not strongly reduce abundance **(Fig. S6B)**. This distinction is especially relevant for PTEN, where highly stable but catalytically inactive variants are known to act as dominant negatives—interfering with the function of the wild-type protein despite retaining normal expression levels. Consistent with this, recurrent PTEN cancer hotspot mutations and known dominant negative variants showed high magnitudes of activity residuals^72^ **(Fig. 4D)**, underscoring the importance of how, not just whether, phosphatase activity is lost.

To assess whether allosteric effects decay with distance from functional sites, we plotted the absolute abundance-corrected activity residuals of each variant against its distance from the bound ligand **(Fig. 4G, 5G)**. In both PTEN and GCK, we observed a clear exponential decay: variants closest to the orthosteric sites had the strongest effects, and the typical effect size dropped by half over distances of 14.5 Å (PTEN) and 15.8 Å (GCK). This distance-dependent pattern was statistically significant in both proteins. Linear regression of the log-transformed absolute residuals against distance yielded negative slopes (PTEN: slope = –0.033, *P* = 2.99e-07; GCK: slope = –0.046, *P* = <2e-016), consistent with a conserved distance-dependent decay of stability-independent functional effects. These findings mirror the distance-dependent decay of mutational effects observed in experimental allosteric maps—including decay from binding interfaces in protein interaction domains^40^,in KRAS^42^, and in activity-altering mutations in the Src kinase^41^, as well as in the allosteric phytohormone receptor PYL1^49^. They are also similar to the decay of abundance corrected mutational effects on predicted fitness presented above **(Fig. 2)**. Together, these results strongly support a conserved biophysical principle of distance-dependent allosteric decay.

Both PTEN and GCK contain clinically pathogenic variants that clustered in the active site and interface regions—as expected—but many were also located at distal positions **(Fig. 4E, 5E)**. To test whether such non-orthosteric pathogenic variants exert functional effects beyond abundance loss, we compared their activity–abundance residuals to those of abundance-matched, non-pathogenic controls. In both orthosteric and non-orthosteric regions, pathogenic variants showed significantly larger residuals than abundance-matched controls **(Fig. 4H, 5H)**, indicating greater loss of activity than expected from abundance effects alone. Many pathogenic variants in PTEN and GCK therefore likely act through allosteric mechanisms.

## Discussion

Taken together, our results indicate that allostery is an important cause of loss-of-function variant pathogenicity. Although many pathogenic variants destabilize proteins^29–33^, changes in stability only partially account for the pathogenicity of variants. Both state-of-the-art computational predictions **(Fig. 1,2)**, and direct experimental measurements **(Fig. 3-5)** indicate that the indirect, allosteric effects of mutations on activity are a frequent mechanism of pathogenicity. In both PTEN and GCK, for example, many known pathogenic variants have no or weak effects on stability but strong experimentally-measured allosteric effects on activity.

To date, comprehensive experimental allosteric maps have only been generated for a handful of proteins^40–42,49,50^ and a limitation of our study is the extrapolation of conclusions from these proteins to the whole proteome. Nevertheless, results from our proteome-wide analyses using state-of-the-art computational predictors **(Fig. 1-2)** are highly consistent with the conclusions from the experimental allosteric maps **(Fig. 3-5)**. Ultimately, it will be important in future work to scale up the production of experimental allosteric maps, particularly for proteins that harbor large numbers of clinically observed variants^38,62,63^. Furthermore, we have only experimentally considered protein-protein binding and enzyme catalytic activity in this study; future efforts should develop methods to quantify allosteric effects on additional molecular activities, such as DNA binding and small-molecule interactions. A comprehensive understanding of pathogenic mechanisms will ultimately require experimental tools capable of capturing allosteric regulation across the full spectrum of disease-relevant molecular activities.

Additional caveats should be noted. First, ESM1v is a proxy for variant pathogenicity and evolutionary fitness derived from protein sequences; while it captures many aspects of mutational impact on protein function, it is not a direct measure of clinical pathogenicity. Second, our computational analyses and the equivalent experimental data aim to quantify allostery but they do not identify the underlying causal molecular mechanisms, which likely involve changes in both conformation and protein dynamics^3,8,73–75^.

However, despite this diversity of possible mechanisms, a conserved principle of allosteric communication appears to be the distance-dependent decay that we have highlighted here. An approximately exponential distance-dependent decay of changes in activity after correcting for abundance changes is seen in all seven experimental allosteric maps published to date^40–42,49,50^ and it is also observed in the PTEN and GCK allosteric maps that we have presented here **(Fig. 4-5)**. It is also visible when plotting computational predictions of variant pathogenicity after accounting for predicted stability effects **(Fig. 2)**. Exponential distance-dependent decay has been observed for evolutionary conservation in enzyme active sites, for chemical shift perturbations in nuclear magnetic resonance (NMR) experiments and for energetic couplings in molecular dynamic simulations^57,58,76^. We believe, therefore, that distance-dependent allosteric decay is likely to be a conserved general principle of protein biophysics.

Pervasive and distance-dependent allostery has important implications for human genetics. First, many loss-of-function variants are likely to have allosteric mechanisms of action. Second, our data predict that pathogenic variants are more likely to have allosteric mechanisms when they are located structurally close to active sites. Third, and conversely, drugs targeting non-orthosteric protein pockets are more likely to have strong allosteric effects when they target pockets closer to a protein’s active site. In future work, it will be important to further refine the rules of allostery, allowing improved mechanistic interpretation of variant pathogenicity and prediction of the most effective sites to target to therapeutically modulate protein function.

## Methods

### Data processing for human proteome-wide predictions

Proteome-wide ESM1v scores were obtained from Zenodo (May 2025)^77^, with detailed methods described in the accompanying publication^78^. AlphaFold predicted structures were downloaded from the AlphaFold Protein Structure Database (June 2025)^79^ and filtered based on criteria established in reference 29^29^, excluding proteins shorter than 16 amino acids or longer than 2700 amino acids. This yielded a final dataset of 20,144 unique, unfragmented human proteins, of which 19,804 had matching ESM1v scores and were retained for downstream analyses.

Stability changes upon mutation (ΔΔG) were predicted for all possible missense mutations using ThermoMPNN v1.0.0^52^ with default settings applied to the AlphaFold-predicted structures.

Protein core positions were identified based on relative solvent-accessible surface area (rSASA), calculated using DSSP with BioPython with default settings^80^. Residues were classified as “core” if they were not predicted to be disordered or part of a binding site and had rSASA < 0.2, following criteria from previous publications^81,82^. Binding and active sites were annotated using proteome-wide predictions from the PIONEER interactome^81^.

Proteins were grouped into functional classes according to the “Function and compartment based” classification provided by The Human Protein Atlas^83–85^, available at https://www.proteinatlas.org/humanproteome/proteinclasses.

Clinical variant annotations were obtained from the supplementary materials of reference 29^29^. Variants were originally extracted from ClinVar^53,54,86^ and processed according to established protocols^31^, retaining only single-nucleotide variants with at least a one-star review status and mapped to GRCh38. For this study, variants labeled pathogenic or likely pathogenic were grouped as “pathogenic”, while those labeled benign or likely benign were grouped as “benign”. Variants with other labels or synonymous mutations were excluded. This yielded 71,418 benign and 42,579 pathogenic clinically annotated variants across 12,336 human proteins. ThermoMPNN-predicted missense mutations were classified as: destabilizing if ΔΔG > 1 kcal/mol, stabilizing if ΔΔG < –0.5 kcal/mol, following the thresholds defined in reference 52^52^.

A subset of allosteric proteins, listed in **Supplemental Table 3**, were selected for focused analyses. These included all full-length human proteins present in two curated datasets: six proteins from the 20-protein benchmark set used to validate the allosteric network model Ohm^87^, and one additional protein, isocitrate dehydrogenase 1 (IDH1), included from reference 88^88^. The enzyme prothrombin was excluded because its allosteric regulator functions as an activator rather than an inhibitor. For glucokinase (GCK), annotations of allosteric sites were based on literature reports rather than crystal structure contacts, as its allosteric regulator (MRK) is also an activator. Orthosteric and allosteric residues were defined based on direct atomic contacts with bound substrates or modulators as observed in crystal structures. Residues were considered in contact if their atoms exhibited van der Waals overlap with atoms of the ligand, as determined using the ChimeraX VDW overlap method^89^.

### Data processing for Mega-scale, Domainome 1.0, and VAMP-seq datasets

ESM1v scores for variants in the Mega-scale dataset were obtained from the ProteinGym database^90^. Of the 331 natural protein domains studied, 64 had corresponding ESM1v predictions available. For the Domainome 1.0 dataset, ESM1v scores were extracted from the supplementary materials of the original publication^33^. ESM1v scores for VAMP-seq variants were derived from the proteome-wide ESM1v dataset described above. VAMP-seq abundance fitness data were collected for nine full-length human proteins: PRKN, OCT, CYP2C9, TPMT, ASPA, NUDT15, VKOR, PTEN, and GCK^38,61,91–96^.

Clinical variant annotations for both the Domainome 1.0 and VAMP-seq proteins were obtained from the supplementary data of reference 29^29^, following the same filtering criteria as described above.

### Variant Classification

- Domainome 1.0 variants were classified based on criteria defined in the original publication. Mutations were considered stabilizing if they had a statistically significant stabilizing effect (*FDR_stabilizing* < 0.1) and a scaled fitness > 0.3. Conversely, mutations were classified as destabilizing if they had a significant destabilizing effect (*FDR_destabilizing* < 0.1) and a scaled fitness < 0.
- VAMP-seq variants were categorized based on abundance thresholds defined by ProteinGym^90^. Mutations falling below the defined abundance cutoff were considered destabilizing.
- Mega-scale variants were classified using thresholds established in the original study ^32^. Note that the ΔΔG directionality in this dataset is inverted relative to others: mutations with ΔΔG > 1 kcal/mol were classified as stabilizing, while those with ΔΔG < –1 kcal/mol were classified as destabilizing.

### Data processing for experimental allosteric maps

Analyses for PSD-95-PDZ3 and GRB2-SH3 were performed using variants with corresponding ESM1v scores, obtained from the human proteome-wide prediction dataset. Orthosteric sites were defined based on published structural annotations as residues within 5 Å of bound ligands^40^. Variants were classified as allosteric if they occurred outside orthosteric sites and exhibited binding ΔΔG (ΔΔGb) values greater than or equal to the mean ΔΔGb of orthosteric-site mutations.

For PTEN and GCK, analyses were based on variants with both abundance and activity fitness measurements. In PTEN, orthosteric site annotations were manually curated from the literature^97–103^. These included the catalytic loops (arginine loop, residues 35–49; WPD loop, 88–98; P loop, 123–130; TI loop, 159–171), calcium-binding regions (residues 200–212, 226–238, and 258–268), the Cα2 loop (residues 327–335), the membrane-binding helix (residues 151–174), and the PI(4,5)P₂-binding motif (PBM, residues 1–15).

In GCK, orthosteric sites were defined based on annotated ligand-contacting residues described in previous structural studies^104–107^. These included glucose-binding residues (positions 151–153, 168–169, 204–206, 225–231, 254–258, 287, and 290) and ATP-binding residues (positions 78–85, 295–296, 331–333, 336, and 410–416).

Variants from gnomAD and ClinVar were downloaded from their respective online portals in April 2025^86,108^. Somatic variant annotations were obtained from the supplementary materials of a previous publication and filtered to retain only cancer hotspot mutations^72^. These hotspots were defined using the “curated set of non-redundant studies” available on cBioPortal^109,110^.

### Statistical analysis and data visualization

Linear regression models were used to analyze proteome-wide variant effect predictions, large-scale abundance datasets, and individual protein domains with experimentally mapped allosteric sites. To evaluate the relative contributions of folding ΔΔG and binding ΔΔG to ESM1v scores, both ordinary linear regression and linear mixed-effects models were employed. Linear regression was performed using the `lm()` function from the stats package in R, while mixed-effects models were fitted using the `lmer()` function from the lme4 package^111,112^. As results from both approaches were highly consistent **(Supplemental Tables 4–7),** linear regression outcomes are reported for interpretability. Adjusted R^2^ values were extracted directly from the `summary()`output of the fitted linear models in R. To estimate variability and provide robust confidence intervals, adjusted R^2^ was also bootstrapped using 1,000 resampling iterations. Model diagnostics included residual and prediction analysis, and model comparisons were performed using ANOVA, with F-statistics and p-values reported.

LOESS residual analyses were performed using only mutations located outside orthosteric sites. LOESS fitting was conducted with the `loess()` function in R, using a span of 0.7 and the family = “symmetric” setting. Distances to ligand-binding sites were computed based on Cα atom coordinates for all proteins, except for PSD-95-PDZ3 and GRB2-SH3, where distances were measured using side chain heavy atoms.

To generate abundance-matched control distributions for residual activity, we performed bootstrap resampling from non-pathogenic variants. For each protein (PTEN and GCK), pathogenic variants with measured abundance and residual activity scores were first binned by their abundance values using fixed-width bins (bin size = 0.5). For each pathogenic variant, we identified all non-pathogenic variants in the same abundance bin (excluding any known pathogenic variants). From these bin-matched pools, one variant was sampled randomly with replacement for each pathogenic variant, and the median residual activity across the matched sample was recorded. This process was repeated 1000 times to generate a distribution of bootstrapped median residuals. The resulting distributions were compared using a two-sided Wilcoxon rank-sum test. Violin plots show the distribution of residuals in pathogenic variants and the distribution of bootstrap medians from abundance-matched controls.

To quantify the distance-dependent decay of mutational effects, nonlinear least-squares regression was applied to the LOESS residuals. The allosteric decay model included two parameters: the decay rate (−b) and the x-intercept (a). This fitting was performed on residuals from either ESM1v versus folding ΔΔG (ΔΔGf) or activity versus abundance fitness datasets, using only mutations with negative residuals that fell below the LOESS curve. Nonlinear regression was conducted with the `nls()` function in R, using initial parameter values of a = 1 and b = 0.1. 95% confidence intervals were calculated by bootstrapping the model parameter with 1,000 replicates.

To identify residues with significantly stronger mutational impacts than predicted by the distance-dependent decay model, we computed residuals between observed and expected values per residue and applied a one-sided Wilcoxon signed-rank test (alternative = “greater”). P-values were adjusted using the Benjamini-Hochberg FDR method. Residues with adjusted *p* < 0.05 were defined as allosteric “hotspots.”

All statistical analyses were performed in R^112^. Experimental schematics were created using <u>BioRender.com</u>, while all other plots were generated using the ggplot2 package^113^. Three-dimensional protein structure visualizations were rendered using ChimeraX v1.6.1^89^.

## Acknowledgment

We thank Taylor Mighell, Mike Thompson, Maximilian R. Stammnitz, Albert Escobedo, Taraneh Zarin, Antoni Beltran, Xianghua Li, and other former and current members of the Lehner lab for feedback and discussions. We also thank the Sanger HumGen/GenGen Informatics for their assistance with the computational infrastructure.

Work in the lab of B.L. was funded by Wellcome (Grant reference: 220540/Z/20/A, ’Wellcome Sanger Institute Quinquennial Review 2021-2026’), a European Research Council (ERC) Advanced (883742) grant, the Spanish Ministry of Science and Innovation (LCF/PR/HR21/52410004, EMBL Partnership, Severo Ochoa Centre of Excellence), Agència de Gestió d’Ajuts Universitaris i de Recerca (AGAUR, 2021 SGR 01226), and the CERCA Program/Generalitat de Catalunya. X.L is funded by the Wellcome Sanger Institute PhD Graduate program.

## Data availability

Proteome-wide predictions of protein stability changes and corresponding ESM1v scores are provided in Supplemental Table 1 (available on Zenodo: https://zenodo.org/uploads/15681969). Raw ThermoMPNN predictions of protein stability changes can be accessed separately at https://zenodo.org/uploads/15586553. Large-scale experimental measurements of protein abundance changes across human domains, including datasets from the Mega-scale and Domainome studies, are accessible via their respective supplementary materials. VAMP-seq abundance data can be obtained from MaveDB and ProteinGym^90,114^. Experimental ΔΔG values for PSD95-PDZ3 and GRB2-SH3 domains are publicly available, with full methodologies described in previous publications^40,115,116^. Experimental measurements of abundance and activity-based fitness for PTEN and GCK can be accessed through the original studies^38,61–63^.

## Code availability

All code and metadata to reproduce the analyses are hosted on GitHub: https://github.com/lehner-lab/allostery_pathogenicity.

## Author contributions

X.L. performed all analyses. X.L. and B.L. conceived the project, designed analyses, and wrote the manuscript.

## Declaration of interests

B.L. is a founder and shareholder of ALLOX.

---

**Fig. S1:**
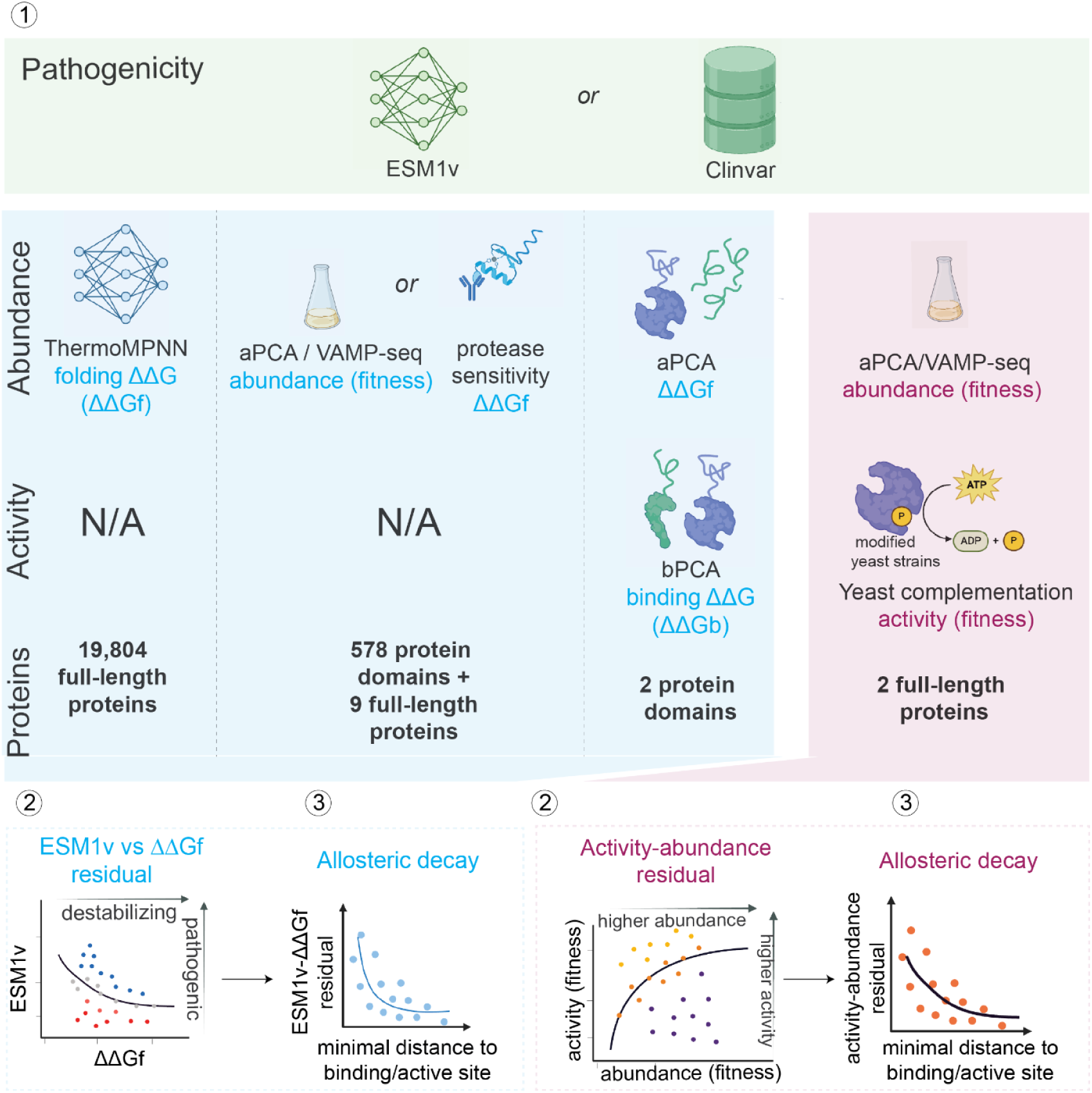
Overview of data types, functional assays, and residual-based analysis strategies. Top panel (green): Summary of protein-scale data used in this study. Pathogenicity scores were derived from either ESM1v or ClinVar annotations. **Middle panel (blue):** Protein abundance was measured using ThermoMPNN-predicted folding free energy changes (ΔΔGf), aPCA/VAMP-seq-based fitness assays, or protease sensitivity. Binding free energy changes (ΔΔGb) was measured using bPCA in two protein domains. **Middle panel (magenta):** Activity was quantified using yeast complementation assays for two full-length proteins, with abundance measured in parallel. **Bottom left (blue):** Schematic of ESM1v–ΔΔGf residual analysis. Residuals quantify the divergence between predicted pathogenicity and measured folding effects. Residuals are then related to distance from the active or binding site to assess allosteric decay. **Bottom right (magenta):** Schematic of activity-abundance residual analysis. The difference between measured activity and abundance (fitness) reflects functional disruption beyond stability effects. These residuals were also regressed against structural distance to active/binding sites to evaluate distance-dependent allosteric effects.

**Fig. S2:**
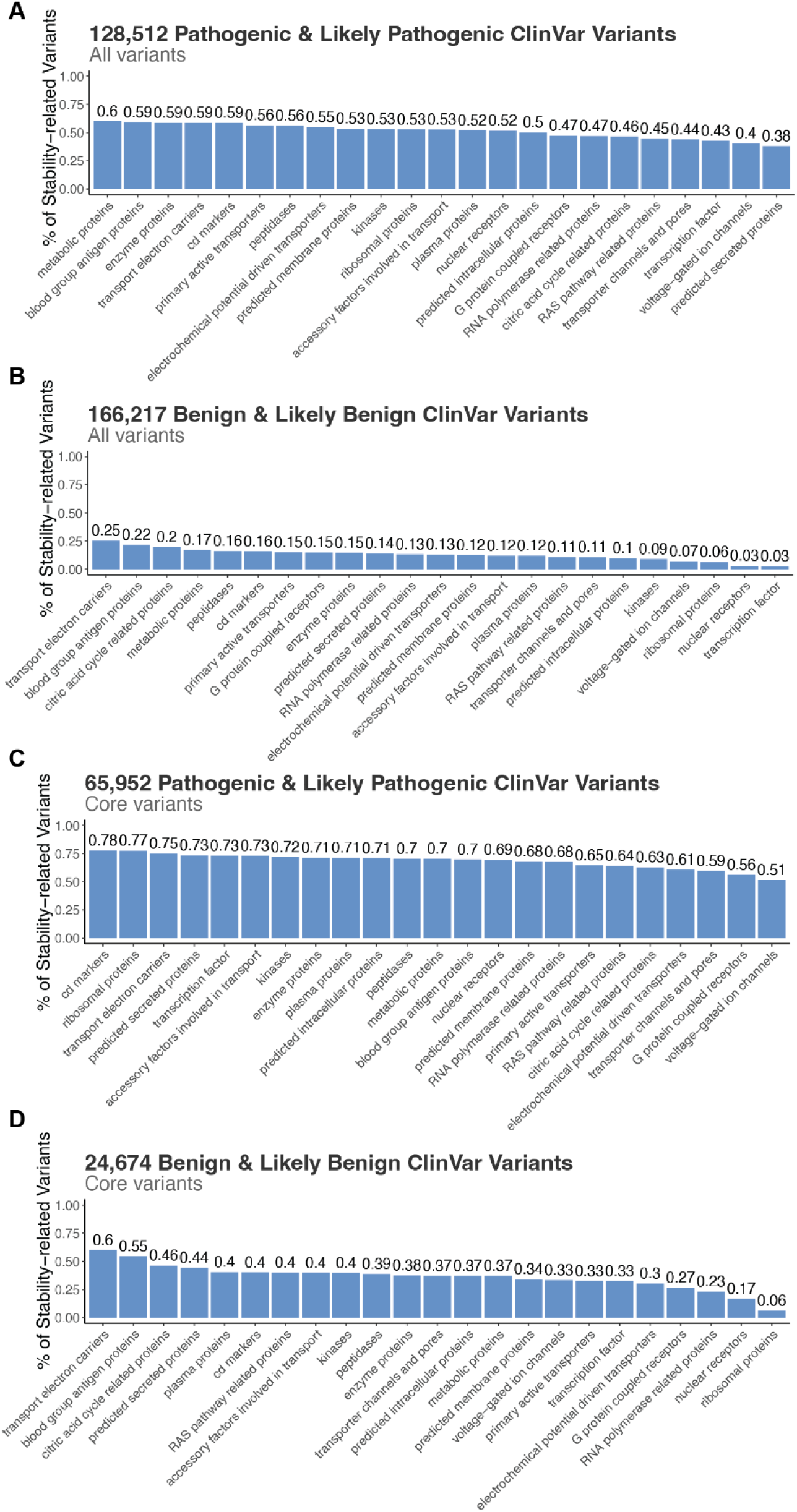
Protein class-specific contribution of stability to pathogenicity. **A**. Bar plot showing the proportion of pathogenic ClinVar variants that are classified as stability-related (destabilizing or stabilizing), across protein functional categories. **B**. Bar plot showing the proportion of benign ClinVar variants that are classified as stability-related, across protein functional categories. **C**. Bar plot showing the proportion of pathogenic ClinVar variants in protein cores (excluding binding sites) that are classified as stability-related, across protein functional categories. **D**. Bar plot showing the proportion of benign ClinVar variants in protein cores (excluding binding sites) that are classified as stability-related, across protein functional categories.

**Fig. S3:**
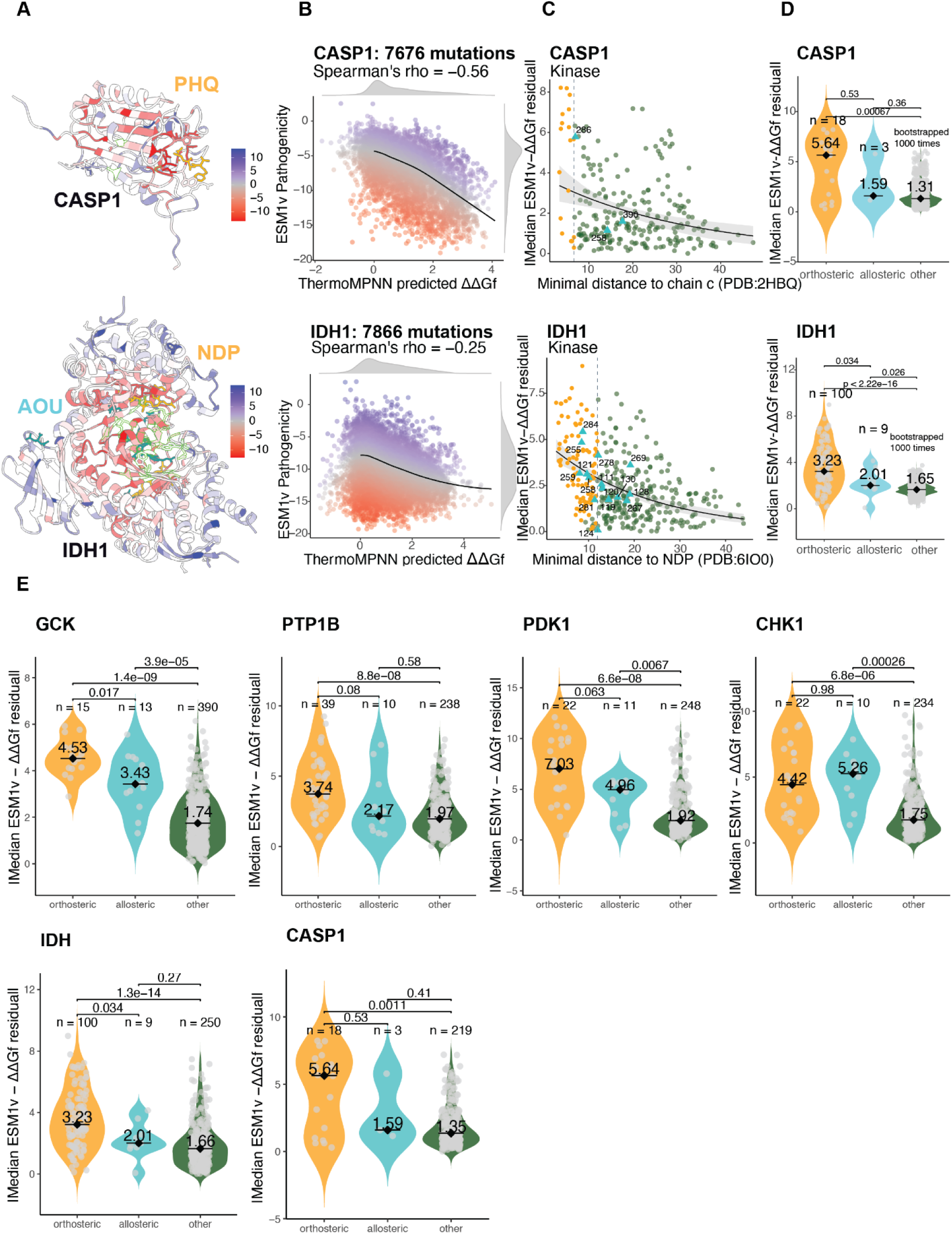
Distance-dependent allosteric decay across diverse protein types. **A**. Crystal structures of CASP1 (PDB: 2FQQ^122^) and IDH1 (PDB: 6IO0^123^), colored by the median residuals between ESM1v and ThermoMPNN (TMPNN) predicted folding free energy changes (ΔΔGf) values. Bound substrates — PHD and NDP — are shown in orange. Allosteric inhibitor AOU is labeled in blue. Annotated allosteric positions are highlighted in neon green. **B**. Scatter plots of ThermoMPNN-predicted ΔΔGf versus ESM1v scores for missense mutations in each protein. A LOESS curve (black line) models the trend, with residuals quantifying deviation between folding impact and predicted pathogenicity. Stable mutations cluster around ΔΔGf ≈ 0. **C**. Per-residue relationship between the absolute median ESM1v–ΔΔGf residual and the minimal Cα distance to the substrates for each structure. Residuals are calculated using only mutations in non-orthosteric sites with negative residuals. An exponential decay model (black curve) was fitted to the data, with 95% confidence intervals shown as grey shading. The grey dashed line indicates the maximum of the minimal distances from any substrate-contacting (orthosteric) residue. Sites are colored by functional class: orthosteric sites in orange circles, non-orthosteric sites in dark green circles, and known allosteric sites (i.e., sites in physical contact with known allosteric regulators) in cyan triangles. **D**. Distributions of the absolute median residuals between ESM1v and ΔΔGf per variant group. For the ‘other’ group, distributions are derived from 1,000 bootstrap replicates, each sampling the same number of non-orthosteric and non-allosteric sites as in the orthosteric group. Violin plots show the distribution shape; black bars and diamonds indicate medians. Sample sizes and median values are annotated. Statistical significance between groups was assessed using two-sided Wilcoxon rank-sum tests. **E**. Distributions of the absolute median residuals between ESM1v and ΔΔGf per variant group across all six allosteric human proteins. The ‘other’ group represents the absolute residuals from all other non-orthosteric and non-allosteric sites. Significance values were calculated using the two-sided Wilcoxon rank-sum test.

**Fig S4:**
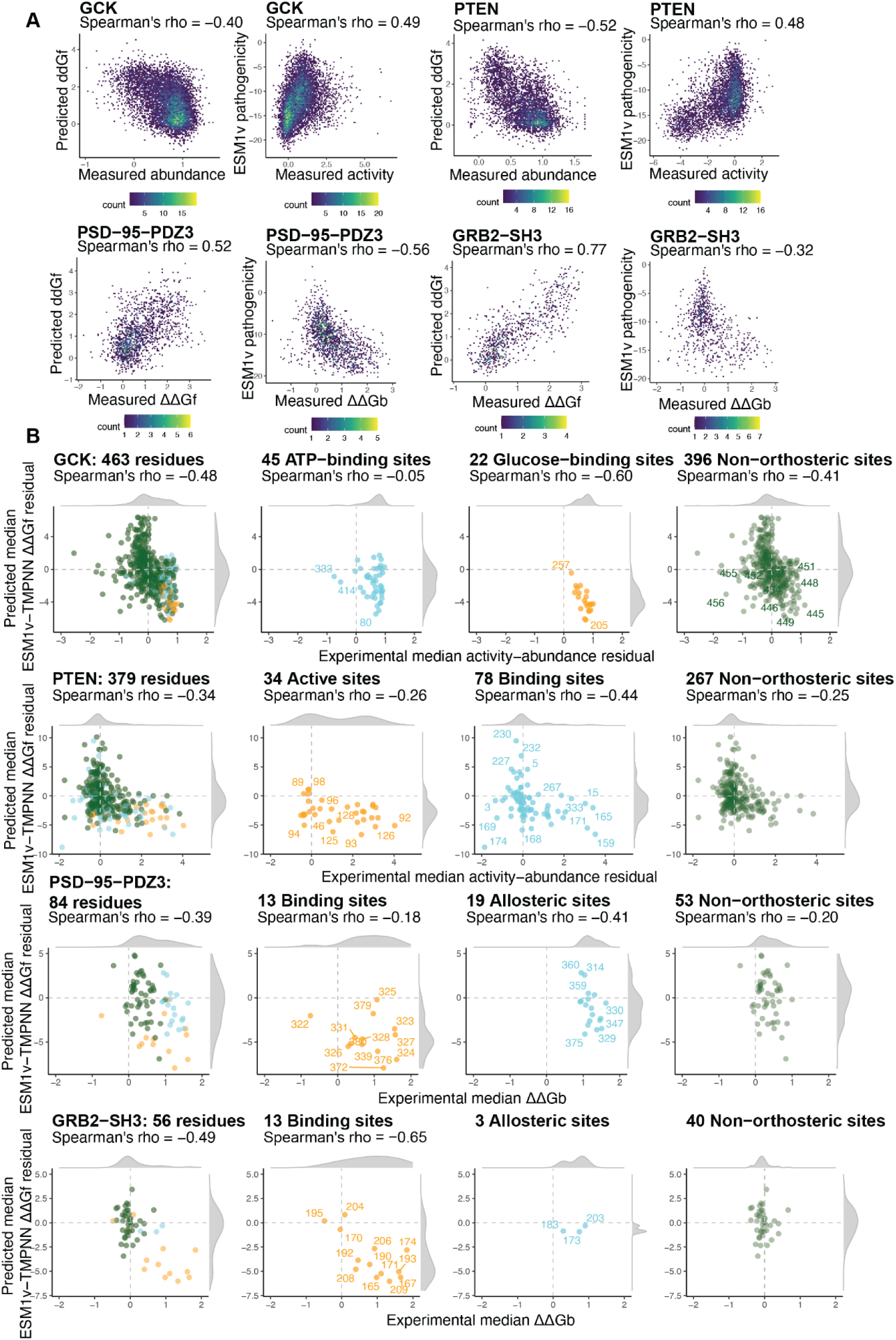
Concordance between experimental and predicted residuals. **A**. Density scatter plots showing the relationship between (left) ThermoMPNN-predicted ΔΔGf and experimentally measured abundance, and (right) ESM1v pathogenicity scores and experimentally measured activity for missense variants in GCK, PTEN, PSD-95-PDZ3, and GRB2-SH3. Point density is color-coded, and Spearman correlation coefficients (ρ) are shown for each plot. **B**. Per-residue comparison between experimentally derived activity–abundance residuals and computationally derived ESM1v–ΔΔGf residuals in GCK, PTEN, PSD95-PDZ3, and GRB2-SH3. Each point represents a single residue, colored by site class. Significant spearman correlations are indicated in each panel.

**Fig. S5:**
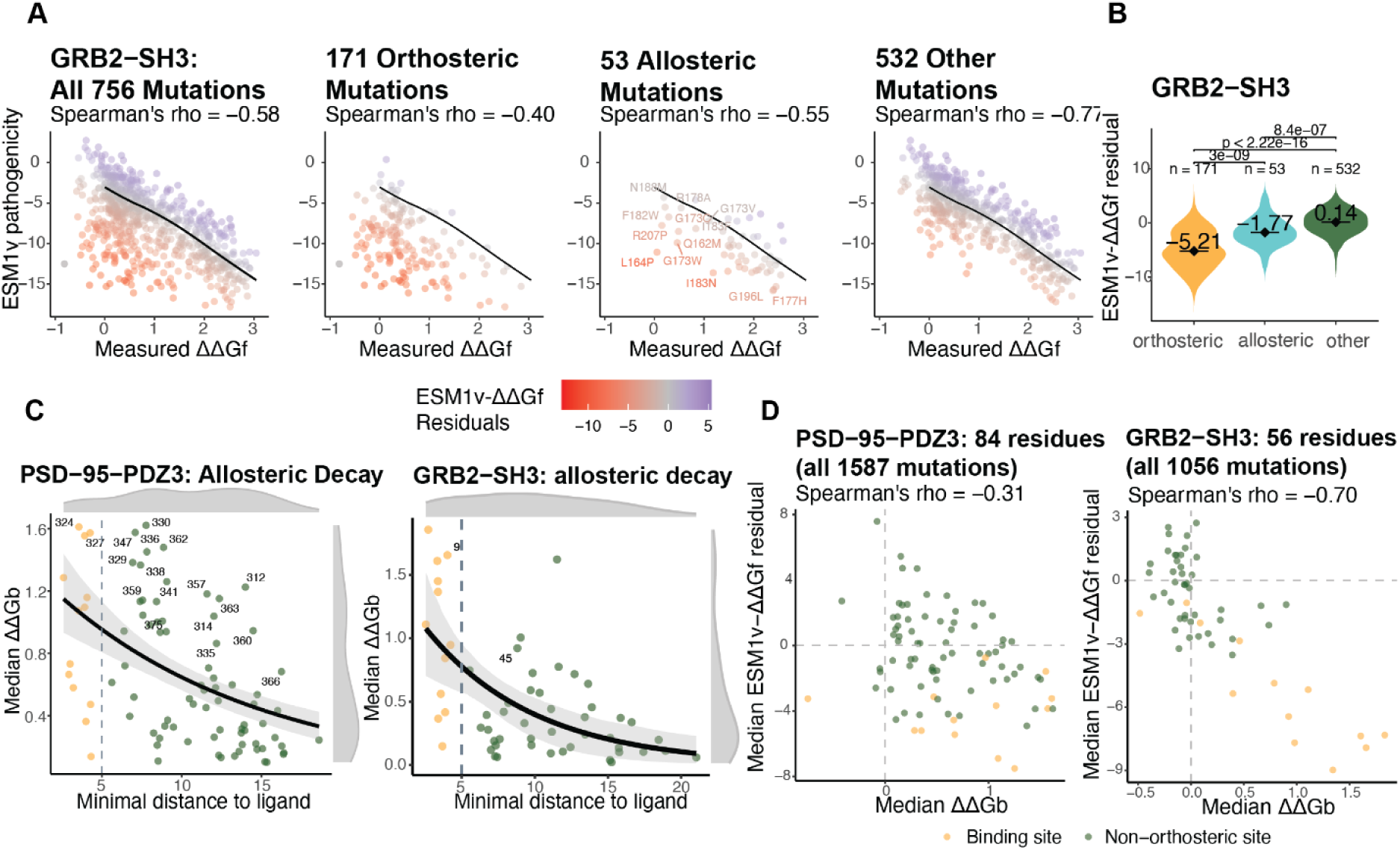
Allosteric decay of binding effects. **A**. Scatter plots showing the relationship between ESM1v pathogenicity scores and experimentally measured folding ΔΔGf for all 756 mutations in GRB2–SH3, with subsets shown for orthosteric (n = 171), allosteric (n = 53), and other (n = 532) mutation classes. Each point is colored by the ESM1v–ΔΔGf residual, and a LOESS curve (black line) models the overall trend. Example allosteric mutations are labeled. **B**. Distributions of the ESM1v–ΔΔGf residuals per variant group. Significance values were calculated using the two-sided Wilcoxon rank-sum test. **C**. Per-residue analysis of binding free energy changes (ΔΔGb) as a function of minimal side-chain distance to the ligand for PSD-95 PDZ3 (left) and GRB2–SH3 (right). Only mutations with ΔΔGb ≥ 0 were included. An exponential decay model was fitted to the data, with 95% bootstrap confidence intervals shown as grey shading. The grey dashed line marks the maximal minimal distance of orthosteric residues (i.e. the outer bound of direct ligand contacts). Residues are color-coded: orthosteric (orange) and non-orthosteric (dark green). Residues with significantly greater ΔΔGb than expected under the decay model (FDR-adjusted *p* < 0.05, one-sided Wilcoxon signed-rank test) are labeled by position number. **D**. Comparison of per-residue median ESM1v–ΔΔGf residuals with median ΔΔGb for PSD-95 PDZ3 (left, 84 residues) and GRB2–SH3 (right, 56 residues).

**Fig. S6:**
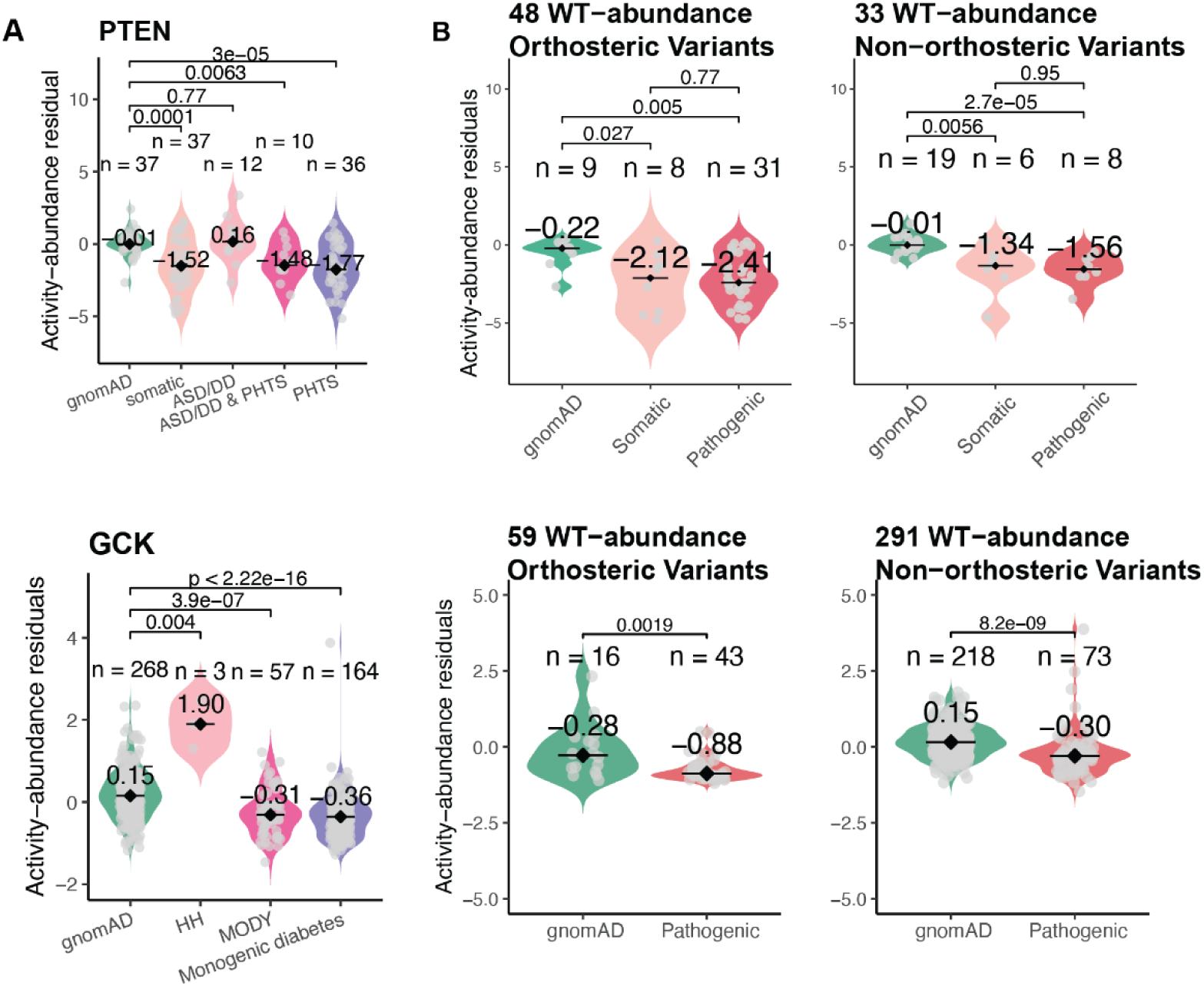
PTEN and GCK analyses extended. **A**. Violin plots showing activity–abundance residuals across variant groups in PTEN (top) and GCK (bottom): gnomAD, and different ClinVar pathogenic categories. Residuals quantify deviation from the fitted activity–abundance relationship. **B**. Violin plots comparing activity–abundance residuals among variants with wild-type–like abundance in PTEN (top) and GCK (bottom), stratified by site type: orthosteric (left) and non-orthosteric (right).

**Supplemental Table 2.**

| Protein | PDB | Function | Name | No. residues with matching ESM1v scores | No. mutations with matching ESM1v scores | References |
| --- | --- | --- | --- | --- | --- | --- |
| GCK | P35557 | Enzyme | Glucokinase | 463 | 8396 | Gersing, S., Schulze, T. K., Cagiada, M., Stein, A., Roth, F. P., Lindorff-Larsen, K., & Hartmann-Petersen, R. (2024). Characterizing glucokinase variant mechanisms using a multiplexed abundance assay. *Genome biology*, 25(1), 98. |
| PTEN | P60484 | Enzyme | Phosphatase and tensin homolog | 383 | 5083 | Matreyek, K. A., Starita, L. M., Stephany, J. J., Martin, B., Chiasson, M. A., Gray, V. E., ... & Fowler, D. M. (2018). Multiplex assessment of protein variant abundance by massively parallel sequencing. *Nature genetics*, 50(6), 874-882. |
| VKOR | Q9BQB6 | Enzyme | Vitamin K epoxide reductase | 162 | 2695 | Chiasson, M. A., Rollins, N. J., Stephany, J. J., Sitko, K. A., Matreyek, K. A., Verby, M., ... & Fowler, D. M. (2020). Multiplexed measurement of variant abundance and activity reveals VKOR topology, active site and human variant impact. *elife*, 9, e58026. |
| NUDT15 | Q9NV35 | Enzyme | Nucleotide triphosphate diphosphatase | 163 | 2922 | Suiter, C. C., Moriyama, T., Matreyek, K. A., Yang, W., Scaletti, E. R., Nishii, R., ... & Yang, J. J. (2020). Massively parallel variant characterization identifies NUDT15 alleles associated with thiopurine toxicity. *Proceedings of the National Academy of Sciences*, 117(10), 5394-5401. |
| ASPA | P45381 | Enzyme | Aspartoacylase | 310 | 5843 | Grønbaek-Thygesen, M., Voutsinos, V., Johansson, K. E., Schulze, T. K., Cagiada, M., Pedersen, L., ... & Hartmann-Petersen, R. (2024). Deep mutational scanning reveals a correlation between degradation and toxicity of thousands of aspartoacylase variants. *Nature Communications*, 15(1), 4026. |
| TPMT | P51580 | Enzyme | Thiopurine S-methyltransferase | 241 | 3648 | Matreyek, K. A., Starita, L. M., Stephany, J. J., Martin, B., Chiasson, M. A., Gray, V. E., ... & Fowler, D. M. (2018). Multiplex assessment of protein variant abundance by massively parallel sequencing. *Nature genetics*, 50(6), 874-882. |
| CYP2C9 | P11712 | Enzyme | Cytochrome P450 enzyme | 486 | 6370 | Amorosi, C. J., Chiasson, M. A., McDonald, M. G., Wong, L. H., Sitko, K. A., Boyle, G., ... & Dunham, M. J. (2021). Massively parallel characterization of CYP2C9 variant enzyme activity and abundance. *The American Journal of Human Genetics*, 108(9), 1735-1751. |
| OCT | O15245 | Transporter | Organic cation transporter 1 | 547 | 9803 | Yee, S. W., Macdonald, C. B., Mitrovic, D., Zhou, X., Koleske, M. L., Yang, J., ... & Coyote-Maestas, W. (2024). The full spectrum of SLC22 OCT1 mutations illuminates the bridge between drug transporter biophysics and pharmacogenomics. *Molecular cell*, 84(10), 1932-1947. |
| PRKN | O60260 | Enzyme | E3 ubiquitin-protein ligase parkin | 465 | 8756 | Clausen, L., Okarmus, J., Voutsinos, V., Meyer, M., Lindorff-Larsen, K., & Hartmann-Petersen, R. (2024). PRKN-linked familial Parkinson's disease: cellular and molecular mechanisms of disease-linked variants. *Cellular and Molecular Life Sciences*, 81(1), 223. |

**Supplemental Table 3.**
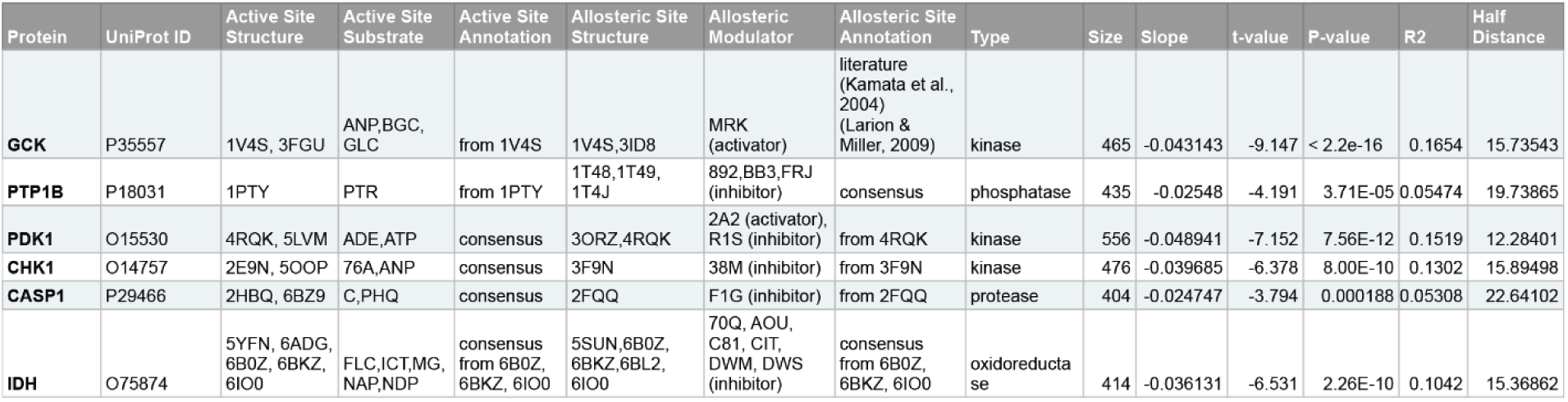

**Supplemental Table 4.** PSD-95-PDZJ: linear regression models.

|  | Model Comparison | Res.Df | RSS | Df | Sum of Sq | F | Pr(>F) |
| --- | --- | --- | --- | --- | --- | --- | --- |
| whole protein | ESM1v ~ $\Delta\Delta G_f$ | 1565 | 21,260 | | | | |
| whole protein | ESM1v ~ $\Delta\Delta G_f + \Delta\Delta G_b$ | 1564 | 18,199 | 1 | 3,061.20 | 263.07 | < 2.2e-16 *** |
| non-orthosteric sites | ESM1v ~ $\Delta\Delta G_f$ | 1323 | 15,721 | | | | |
| non-orthosteric sites | ESM1v ~ $\Delta\Delta G_f + \Delta\Delta G_b$ | 1322 | 13,441 | 1 | 2279.3 | 224.18 | < 2.2e-16 *** |
| orthosteric sites | ESM1v ~ $\Delta\Delta G_f$ | 240 | 1845 | | | | |
| orthosteric sites | ESM1v ~ $\Delta\Delta G_f + \Delta\Delta G_b$ | 239 | 1704.8 | 1 | 140.21 | 19.657 | 1.413e-05 *** |

**Supplemental Table 5.** GRB2-SHJ: linear regression model.

|  | Model Comparison | Res.Df | RSS | Df | Sum of Sq | F | Pr(>F) |
| --- | --- | --- | --- | --- | --- | --- | --- |
| whole protein | ESM1v ~ $\Delta\Delta G_f$ | 754 | 7,968 | | | | |
| whole protein | ESM1v ~ $\Delta\Delta G_f + \Delta\Delta G_b$ | 753 | 5,795 | 1 | 2,173.30 | 282.41 | < 2.2e-16 *** |
| non-orthosteric sites | ESM1v ~ $\Delta\Delta G_f$ | 583 | 3,265 | | | | |
| non-orthosteric sites | ESM1v ~ $\Delta\Delta G_f + \Delta\Delta G_b$ | 582 | 3,020 | 1 | 245.48 | 47.311 | 1.568e-11 *** |
| orthosteric sites | ESM1v ~ $\Delta\Delta G_f$ | 169 | 1864.3 | | | | |
| orthosteric sites | ESM1v ~ $\Delta\Delta G_f + \Delta\Delta G_b$ | 168 | 1343 | 1 | 521.27 | 65.205 | 1.245e-13 *** |

**Supplemental Table 6.** PSD-95-PDZ3: linear mixed models.

|  | Model Comparison | npar | AIC | BIC | logLik | -2logLik | Chisq | Df | Pr(>Chisq) |
| --- | --- | --- | --- | --- | --- | --- | --- | --- | --- |
| whole protein | ESM1v ~ $\Delta\Delta G_f + (1 \vert \text{pos})$ | 4 | 7575.2 | 7596.6 | -3783.6 | 7567.2 | | | |
| whole protein | ESM1v ~ $\Delta\Delta G_f + \Delta\Delta G_b + (1 \vert \text{pos})$ | 5 | 7409.7 | 7436.5 | -3699.9 | 7399.7 | 167.48 | 1 | < 2.2e-16 *** |
| non-orthosteric sites | ESM1v ~ $\Delta\Delta G_f + (1 \vert \text{pos})$ | 4 | 6354.3 | 6375.1 | -3173.2 | 6346.3 | | | |
| non-orthosteric sites | ESM1v ~ $\Delta\Delta G_f + \Delta\Delta G_b + (1 \vert \text{pos})$ | 5 | 6215.8 | 6241.7 | -3102.9 | 6205.8 | 140.57 | 1 | < 2.2e-16 *** |
| orthosteric sites | ESM1v ~ $\Delta\Delta G_f + (1 \vert \text{pos})$ | 4 | 1175.3 | 1189.2 | -583.64 | 1167.3 | | | |
| orthosteric sites | ESM1v ~ $\Delta\Delta G_f + \Delta\Delta G_b + (1 \vert \text{pos})$ | 5 | 1155.6 | 1173 | -572.78 | 1145.6 | 21.705 | 1 | 3.179e-06 *** |

**Supplemental Table 7.** GRB2-SH3: linear mixed models.

|  | Model Comparison | npar | AIC | BIC | logLik | -2logLik | Chisq | Df | Pr(>Chisq) |
| --- | --- | --- | --- | --- | --- | --- | --- | --- | --- |
| whole protein | ESM1v ~ $\Delta\Delta G_f + (1 \vert \text{pos})$ | 4 | 3339.9 | 3358.4 | 1666 | 3331.9 | | | |
| whole protein | ESM1v ~ $\Delta\Delta G_f + \Delta\Delta G_b + (1 \vert \text{pos})$ | 5 | 3304.1 | 3327.2 | -1647 | 3294.1 | 37.858 | 1 | 7.609e-10 *** |
| non-orthosteric sites | ESM1v ~ $\Delta\Delta G_f + (1 \vert \text{pos})$ | 4 | 2461.3 | 2478.8 | -1226.7 | 2453.3 | | | |
| non-orthosteric sites | ESM1v ~ $\Delta\Delta G_f + \Delta\Delta G_b + (1 \vert \text{pos})$ | 5 | 2434.5 | 2456.4 | -1212.3 | 2424.5 | 28.795 | 1 | 8.047e-08 *** |
| orthosteric sites | ESM1v ~ $\Delta\Delta G_f + (1 \vert \text{pos})$ | 4 | 810.36 | 822.92 | -401.18 | 802.36 | | | |
| orthosteric sites | ESM1v ~ $\Delta\Delta G_f + \Delta\Delta G_b + (1 \vert \text{pos})$ | 5 | 807.81 | 823.52 | -398.91 | 797.81 | 4.5424 | 1 | 0.03307 * |

## Notes

### Summary of Updates

Reformatted figures to avoid watermark overlap Moved figure legends next to figures Added missing legend for panel S5B Added sentence 'Annotated allosteric positions' to Fig 2B legend Added 'protein cores' label to Fig 1G Corrected 19,803 → 19,804 in Fig S1 Added citations for PDB structures + new reference 16

